# Ephrin-B1 blocks adult cardiomyocyte proliferation and heart regeneration

**DOI:** 10.1101/735571

**Authors:** Marie Cauquil, Céline Mias, Céline Guilbeau-Frugier, Clément Karsenty, Marie-Hélène Seguelas, Gaël Genet, Edith Renaud-Gabardos, Anne-Catherine Prats, Véronique Pons, Maxime Branchereau, Christophe Heymes, Denis Calise, Olivier Lairez, Danièle Daviaud, Benjamin Honton, Céline Frongia, Bernard Ducommun, Marie-Bernadette Delisle, Dina N. Arvanitis, Atul Pathak, Jean-Michel Sénard, Céline Galés

**Affiliations:** Institut des Maladies Métaboliques et Cardiovasculaires, Université de Toulouse, INSERM, Toulouse, France; Département d’Histopathologie, Centre Hospitalier Universitaire de Toulouse, Université de Toulouse, Toulouse, France; Département de Cardiologie, Centre Hospitalier Universitaire de Toulouse, Université de Toulouse, Toulouse, France; Département de Microchirurgie, Université de Toulouse, INSERM, Toulouse, France; INSERM, UMR 1043, Centre de Physiopathologie de Toulouse Purpan, Microscopy platform, Toulouse, France; Département de médecine cardiovasculaire, unité d’hypertension, facteurs de risque insuffisance cardiaque, Clinique Pasteur, Toulouse, France; ITAV, Université de Toulouse, CNRS, Toulouse, France; Département de Pharmacologie Clinique, Centre Hospitalier Universitaire de Toulouse, Université de Toulouse, Toulouse, France

**Keywords:** cardiomyocyte proliferation, cardiac regeneration, cardiomyocyte morphology, myocardial infarction, apectomy, Ephrin-B1, Yap1

## Abstract

**Aims:** Deciphering the innate mechanisms governing the blockade of proliferation in adult cardiomyocytes (CMs) is challenging for mammalian heart regeneration. Despite the exit of CMs from the cell cycle during the postnatal maturation period coincides with their morphological switch to a typical adult rod-shape, whether these two processes are connected is unknown. Here, we examined the role of ephrin-B1, a CM rod-shape stabilizer, in adult CM proliferation and cardiac regeneration.

**Methods and results:** Transgenic- or AAV9-based ephrin-B1 repression in adult mouse heart led to substantial proliferation of resident CMs and tissue regeneration to compensate for apex resection, myocardial infarction (MI) and senescence. Interestingly, in the resting state, CMs lacking ephrin-B1 did not constitutively proliferate, indicative of no major cardiac defects. However, they exhibited proliferation-competent signature, as indicated by higher mononucleated state and a dramatic decrease of miR-195 mitotic blocker, which can be mobilized under neuregulin-1 stimulation *in vitro* and *in vivo*. Mechanistically, the post-mitotic state of the adult CM relies on ephrin-B1 sequestering of inactive phospho-Yap1, the effector of the Hippo-pathway, at the lateral membrane. Hence, ephrin-B1 repression leads to phospho-Yap1 release in the cytosol but CM quiescence at resting state. Upon cardiac stresses (apectomy, MI, senescence), Yap1 could be activated and translocated to the nucleus to induce proliferation-gene expression and consequent CM proliferation

**Conclusions:** Our results identified ephrin-B1 as a new natural locker of adult CM proliferation and emphasize that targeting ephrin-B1 may prove a future promising approach in cardiac regenerative medicine for HF treatment.

**Significance:** The mammalian adult heart is unable to regenerate due to the inability of cardiomyocytes (CMs) to proliferate and replace cardiac tissue lost. Exploiting CM-specific transgenic mice or AAV9-based gene therapy, this works identifies ephrin-B1, a specific rod-shape stabilizer of the adult CM, as a natural padlock of adult CM proliferation for compensatory adaptation to different cardiac stresses (apectomy, MI, senescence), thus emphasizing a new link between the adult CM morphology and their proliferation potential. Moreover, the study demonstrates proof-of-concept that targeting ephrin-B1 may be an innovative therapeutic approach for ischemic heart failure.

## Introduction

In the recent years, reinitiating the proliferative activity of differentiated resident cardiomyocytes (CMs) has emerged as an exciting avenue for cardiac regenerative medicine^1^. Theoretically, this challenging concept could provide the opportunity to yield sufficient CMs to replace their profuse loss after myocardial damage. Several studies have supported the natural self-renewal potential of mammalian adult CMs, but this mechanism has remained too weak to efficiently repair the heart^2-4^. Thus, strategies now focus on boosting this regenerative process by exploiting factors specifically involved in the early CM proliferation arrest during the postnatal period^5-9^, and they have succeeded in extending the neonatal regenerative window. In addition, several studies have recently explored the adult stage and identified factors allowing for adult CM proliferation, as well as heart regeneration, despite low proliferation rates^5, 10-16^. However, in the absence of cardiac injury, these studies did not report the *in vivo* consequences of the modulated factor on CM proliferation at baseline, or showed constitutive CM proliferation, most likely reflecting a functional compensatory mechanism and precluding these factors as *bona fide* targets for future clinical regenerative therapies. Hence, identification of the natural molecular mechanisms governing the postmitotic state of adult CMs remains a crucial issue.

*In situ*, mammalian CMs stop dividing around postnatal day 7 (P7), consistent not only with the P1-P7 regenerative cardiac window in neonatal mice^17^ but also with the extinction of growth factors^10, 16, 18^, CM cell cycle regulators^15, 19^ and cell growth pathways^20^. Interestingly, the exit of CMs from the cell cycle during the postnatal maturation period also coincides with a morphogenesis step during which the CM switches from a round-shape to a mature rod-shape^21^. This polarization process of the CM, reminiscent of the polarity in epithelial cells, results in an asymmetric organization of plasma membrane components underlying specific functions, *i*.*e*. the intercalated disk involved in direct CM-CM longitudinal interactions and the lateral membrane connected to the basement membrane and the extracellular matrix (ECM). Until now, the molecular events leading to the establishment of the CM rod-shape remain unknown. Despite cell polarization and proliferation generally are generally considered as antagonistic processes, whether adult CM morphology and proliferation arrest might be directly connected has never been explored.

Recently, we identified the transmembrane protein ephrin-B1, a member of the large family of eph receptors/ephrin ligands^22^, as a new constituent of the adult CM lateral membrane and acting as a stabilizer of the adult CM rod-shape. We demonstrated that this protein acts independently of the costamere structure at the lateral membrane in agreement with the lack of contractile defects in young adult *efnb1* knockout (KO) mice^23^. Ephrin-B1 does not directly contribute to the establishment of the adult CM rod-shape, *i*.*e*., the asymmetry of the surface membrane. However, it participates in the maturation of the adult CM polarity since we showed that it stabilizes this specific morphology through a physical stretching of the lateral membrane, which is essential for the overall cardiac tissue cohesion in adult mice. Accordingly, mice with global (KO) or CM-specific (cKO) *efnb1* deletion display a soft cardiac tissue with an architectural disorganization of the CM lateral membrane leading to spindle-shaped, immature-like CMs. Interestingly, while 2-month-old *efnb1* KO mice exhibit cardiac tissue disorganization, they do not develop cardiac failure in homeostatic conditions which was quite surprising given the importance of the tissue cohesion in the stabilization of the cardiac function. Nevertheless, they were intolerant to cardiac mechanical stress, in agreement with the lack of tissue cohesion. Here, we now questioned about a potential role for ephrin-B1 in the adult CM proliferation state that could reconcile the loss of the adult CM rod-shape and tissue cohesion to the standard cardiac function in young adult *efnb1*-KO mice.

## Methods

Detailed materials and methods are provided in the Supplementary material online.

### Animal models

Global (KO) and CM-specific (cKO) *efnb1* KO mice have already been described ^23^. *Efnb1* WT and KO mice were kept in a mixed S129/S4 × C57BL/6 background. All studies were performed on male and age-matched mice. Experimental animal protocols were carried out in accordance with the French regulation guidelines for animal experimentation and were approved by the French CEEA-122 ethical committee.

### Data analysis

The *n* number for each experiment and analysis are stated in each figure legend. All bar graphs, except for ***Figure 4C*** and ***Supplementary material online*, *Figure S6*A, D, G, H** represent means ± s.e.m. Statistical analyses were performed using Prism v5 software. * *P*<0.05, ** *P*<0.01, *** *P*<0.001 were considered statistically significant; ns: not statistically significant.

**Figure 1.**
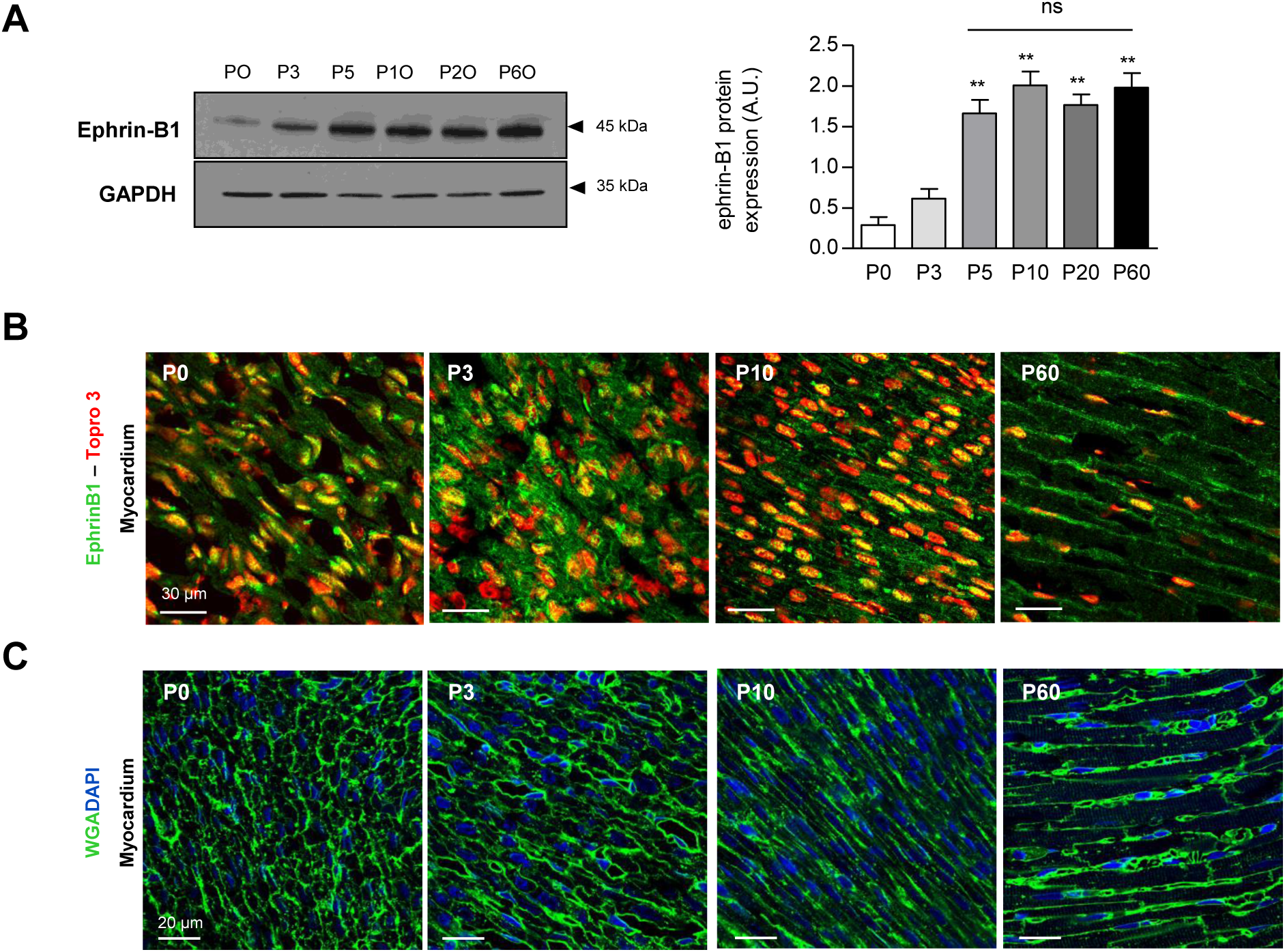
**A**, *left panel*, western-blot analysis of ephrin-B1 in cardiac tissue from rats at different postnatal maturation days (postnatal day 0/P0, day 3/P3, day 10/P10, day 60/P60). Each line represent one rat. *Right panel*, quantification of ephrin-B1 expression normalized to total protein (n=3-5 rats). Data are presented as mean ± SEM and analyzed using one-way ANOVA with Dunett’s post hoc test. ** P<0.01. **B-C**, Ephrin-B1 (**B**) and CM morphology (WGA) (**C**), visualized by immunofluorescence on heart cryosections (myocardium) (**B**) or paraffin-embedded sections (**C**) from rats at different postnatal maturation days (P0, P3, P10, P60). Cell nuclei were immunostained with Topro3 or DAPI markers.

**Figure 2.**
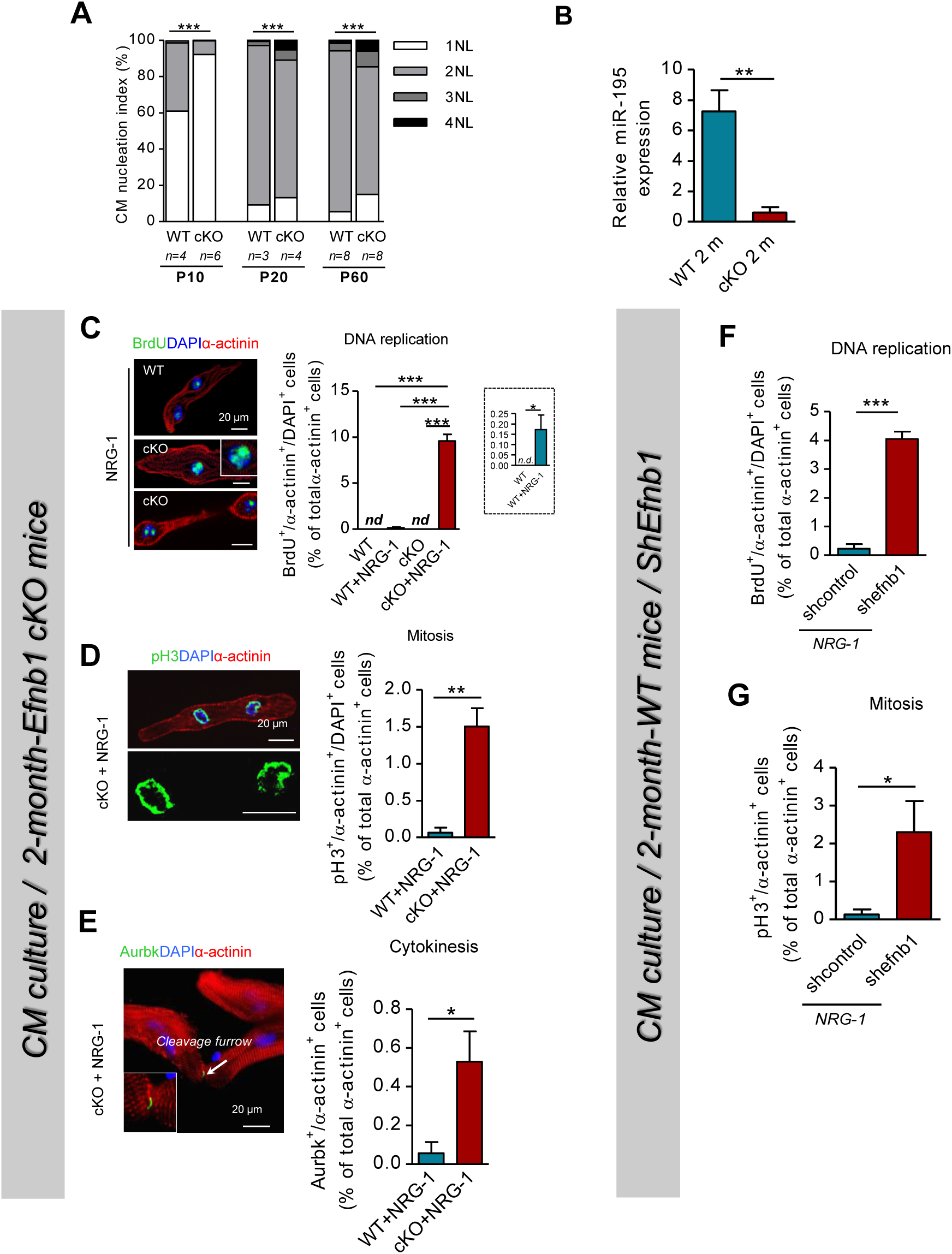
Two-month-old *efnb1 c*KO mice exhibit proliferation-competent CMs *in vitro*. **A**, Nucleation profile of CMs isolated from mice at postnatal days 10, 20 or 60 (P10-P60) (∼350 CMs/mouse). **B**, qRT-PCR quantification of miR-195 expression in CMs isolated from 2-month-old WT and cKO mice (n=4 to 5 mice/group). **C-E**, Replication (BrdU^+^/α-actinin^+^/DAPI^+^), mitosis (pH3^+^/α-actinin^+^/DAPI^+^) and cytokinesis (Aurkb^+^ cleavage furrows/α-actinin^+^) quantified on isolated CMs from 2-month-old WT and cKO mice following 8 days of NRG-1 treatment (∼ 380 CMs/mouse, n=3 to 6 mice/group). **F, G**, quantification of (**F**) replication (BrdU^+^/α-actinin^+^/DAPI^+^; ∼130 CMs/mouse) and (**G**) mitosis (pH3^+^/α-actinin^+^/DAPI^+^; ∼ 500 CMs/mouse) following NRG-1 treatment during 8 days of CMs isolated from 2-month-old WT mice and transduced with shcontrol or sh*efnb1* lentivirus (n=3 mice/group). Data are presented as mean ± SEM and analyzed using Student’s t-test for 2 group comparisons or one-way ANOVA with Tukey post-hoc test for 4 group comparisons * *P*<0.05, ** *P*<0.01, *** *P*<0.001. nd, not detectable.

**Figure 3.**
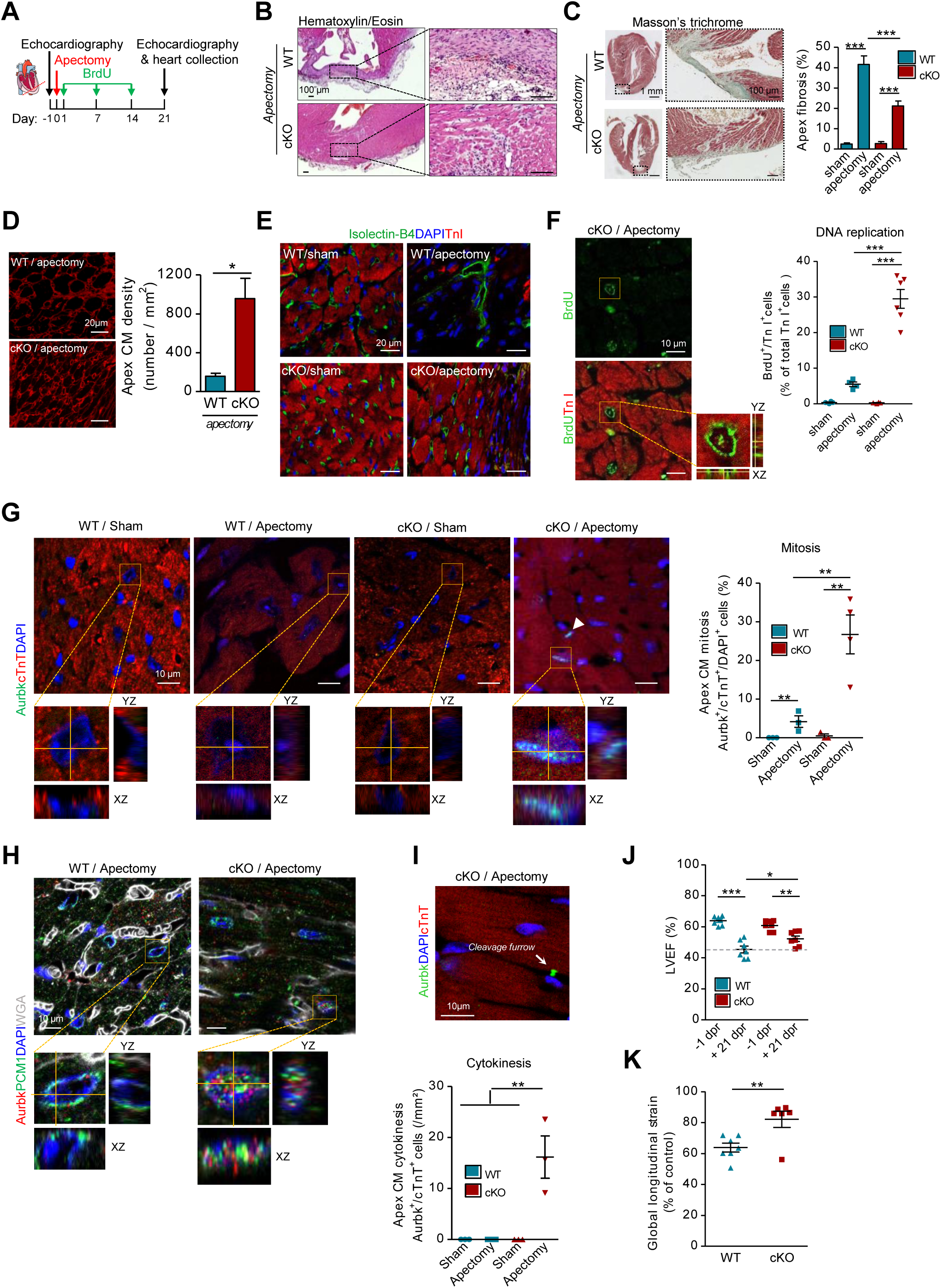
Ephrin-B1 deletion promotes cardiac regeneration following apectomy via adult CM proliferation. **A**, Schematic of apectomy experimental protocol in 2-month-old WT or cKO mice. **B**, Representative hematoxylin/eosin staining of hearts 21 days after apectomy. **C**, Quantification of cardiac fibrosis in the apex (∼3 sections per mice, WT/sham n=4; WT/apec n=6; cKO/sham n=6; cKO/apec n=8). **D**, Quantification of CM density in the apex (∼125 CMs/mouse; WT n=4; cKO n=5). **E**, Representative vascular organization in the apex (isolectin-B4-staining). **F, G**, *In situ* quantification of CMs (**F**) replication (BrdU^+^/TroponinI^+^;KO/apec n=6; n=4 for all others) and (**G**) mitosis (Aurkb^+^/ cTnT^+^/DAPI^+^; cKO/apec n=4, for all others n=3). **H**, Representative images of resident CM mitosis in cardiac tissue from apectomized WT or cKO mice (PCM1^+^/DAPI^+^/Aurkb^+^). **I**, *In situ* quantification of CM cytokinesis (Aurkb^+^ cleavage furrows/cTnT^+^; n=3/group). **J, K**, Echocardiography-based analysis of (**J**) ejection fraction (LVEF, left ventricle ejection fraction; n=7/group) and (**K**) global longitudinal strain (WT n=7; cKO n=6) measured 1 day before (−1 dpr, day post-resection) and 21 days after (+21 dpr) apectomy. Global longitudinal strain represents cardiac deformation at +21 dpr and is expressed as the percentage of strain measured at −1 dpr. Data are presented as mean ± SEM and analyzed using Student’s t-test for 2 group comparisons or one-way ANOVA with Tukey’s post hoc test for 4 group comparisons. * *P*<0.05, ** *P*<0.01, *** *P*<0.001.

**Figure 4.**
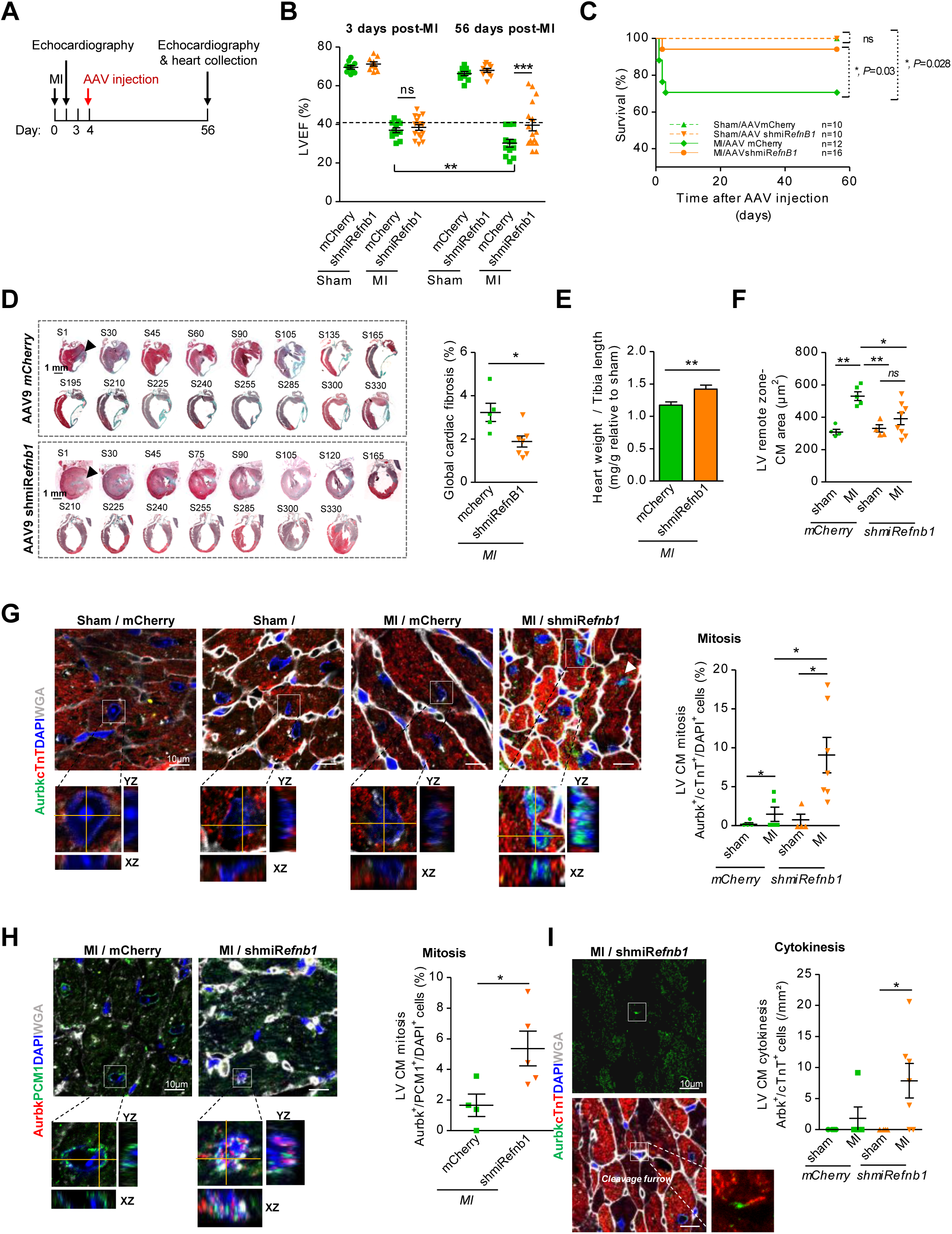
Ephrin-B1 deletion promotes cardiac regeneration following myocardial infarction via adult CM proliferation. **A**, Schematic of the myocardial infarction (MI) protocol using control-(mCherry) or shmiR*efnb1*-AAV9 in 2-month-old mice. **B**, Left ventricle ejection fraction (LVEF) measured by echocardiography 3 or 56 days post-MI (n= 10-16 mice/group). **C**, Kinetics of mouse survival. Groups compared by Chi^2^ Test. **D**, Cardiac fibrosis quantification (24 longitudinal sections, S/heart; MI/mCherry n=5; MI/shmiR*efnb1* n=7). **E**, Heart weight/tibia length ratios (MI/mCherry n=12; MI/shmiR*efnb1* n=16). **F**, *In situ* quantification of CM area (∼50 CMs/mouse; Sham n=4; MI/mCherry n=5; MI/shmiR*efnb1*n=8). **G-I**, *In situ* quantification of CM (**G, H**) mitosis (**G**, Aurkb^+^/DAPI^+^/cTnT^+^ or **H**, PCM1^+^/DAPI^+^/Aurkb^+^). and (**I**) cytokinesis (Aurkb^+^ cleavage furrows/cTnT^+^) in the left ventricle remote zone (Sham n=4; MI/mCherry n=5, MI/shmiR*efnb1* n=7). Data are presented as mean ± SEM and analyzed using Student’s *t*-test for 2 group comparisons, one-way ANOVA with Tukey post-hoc test for 4 group comparisons or two-way ANOVA for echocardiography analysis. * *P*<0.05, ** *P*<0.01, *** *P*<0.001. ns, not significant.

## Results

### Ephrin-B1 postnatal expression correlates with the setting of the CM rod-polarity and the CM proliferation arrest

To investigate whether ephrin-B1 could regulate CM proliferation, we first examined its expression in the cardiac tissue at the postnatal stage during which the CM stop to proliferate. Ephrin-B1 protein expression picked at postnatal day 5 (***Figure 1A***), a time point that correlated with the initiation of ephrin-B1 targeting to the CM lateral membrane (***Figure 1B***) but also with the CM cell cycle withdrawal^24^. Interestingly, the setting of ephrin-B1 at the CM lateral membrane paralleled the onset of the CM adult rod-shape (***Figure 1C***) with the different components structuring the different domains of the polarized plasma membrane, *i*.*e*. the lateral membrane (β-sarcoglycan) with its basement membrane (laminin 2), Supplementary material online, ***Figure S1A***) and the ID (N-Cadherin, Zo-1, Supplementary material online, ***Figure S1B***). At early postnatal stages, ephrin-B1 expression predominated in CM nuclei, which persisted until the adult stage (***Figure 1B***).

### Ephrin-B1 blocks proliferation-competence to adult CMs

We next examined the role of ephrin-B1 in the adult CM proliferation. For that purpose, we first assessed the global CM proliferation in the hearts from 2-month-old adult *efnb1 c*KO mice (CM-specific KO) using immunostaining for Ki67, a marker of all active phases of the cell cycle (G1, S, G2, M). While substantial Ki67-positive CM nuclei were observed in heart tissues from postnatal P2 mice, we did not detect any Ki67 positive staining in the CMs from cKO or WT mice (Supplementary material online, ***Figure S2A***). This result most likely indicated that the absence of ephrin-B1 in the adult CM does not promote constitutive CM proliferation at resting state. Given that both CM proliferation or hypertrophy occur as compensatory mechanisms to meet the increased functional demand of the heart, the lack of constitutive CM proliferation in the absence of ephrin-B1 in young mice hearts is consistent with the lack of functional contractile defect in 2-month-old *efnb1 c*KO mice that we previously reported^23^.

However, the lack of CM proliferation events *in vivo* does not necessarily mean that *efnb1* cKO CMs cannot proliferate. To explore this possibility, we assessed *in vitro* the potential proliferation-competent signature of CMs isolated from 2-month-old *efnb1 c*KO mice. The purity of the isolated CMs was controlled and reached about 80 % in all cases (not shown). We first examined the CM nucleation profile since several studies have demonstrated that mononucleated CMs have a higher proliferation propensity than binucleated one^12^. In agreement, at young adulthood (P60, 2 months old), cKO hearts harbored increased number of mononucleated CMs with, conversely, relatively fewer binucleated CMs but also an increased number of tri- and quadri-nucleated CMs, compared with WT (***Figure 2A***, Supplementary material online, ***Figure S2B***). This defect in CM nucleation from adult cKO mice was even more evident when examining the postnatal cardiac maturation (P10 and P20) during which CMs progressively lose their proliferative aptitude (***Figure 2A***, Supplementary material online, ***Figure S2B***). Moreover, and consistent with a previous report demonstrating the role of miR-195 as a specific mitotic blocker of adult CMs^8^, we also found a dramatic decrease in miR-195 expression in CMs isolated from 2-month-old cKO mice (***Figure 2B***), further reinforcing the proliferative potential of these mice at resting state.

We next determined directly whether the proliferative potential of CMs isolated from young adult *efnb1* cKO could be mobilized upon exposure to neuregulin-1 (NRG-1), a growth factor able to promote adult CM proliferation^11^. *In vitro*, 8-day culture of 2-month-old WT and cKO CMs, in the presence of recombinant human NRG-1, resulted in low but detectable bromodeoxyuridine (BrdU) uptake in WT CMs (0.17±0.07 %), in agreement with a previous report^11^, while the percentage of replicative CMs considerably increased in cKO cells (9.57±0.70 %) (***Figure 2C***). Further supporting CM proliferation, CMs from cKO hearts also exhibited a substantially higher frequency than WT of mitosis (*pH3*^*+*^, ***Figure 2D***) and cytokinesis (*aurora kinase B*^*+*^ *cleavage furrows*, ***Figure 2E***) events. Proliferation signature of NRG-1-treated CMs from *efbn1* cKO mice was reinforced by: **i**. morphological observations showing myofibril delocalization at the CM periphery as previously reported^7, 11, 25^ and additional nuclear rounding and delocalization in numerous cKO CMs (Supplementary material online, ***Figure S2C***), and **ii**. in agreement with Sadek *et al*^26^, epigenetic analysis, showing a majority of CMs with diffuse DNA-euchromatin (Supplementary material online, ***Figure S2D***), consistent with the presence of gene activating-acetylated H3K9/14Ac (Supplementary material online, ***Figure S2E***), in contrast to major DNA-containing foci indicative of heterochromatin in NRG-1-untreated cKO CMs (Supplementary material online, ***Figure S2D***) and correlating with the expression of gene silencing-methylated H3K9me3 (Supplementary material online, ***Figure S2E***). The capacity of adult CMs from *efnb1* cKO mice to proliferate in response to NRG-1 did not arise from CM adaptation that could have occurred during the postnatal maturation, but truly reflected a specific role of ephrin-B1 at the adulthood. Indeed, lentiviral-based *efnb1* deletion but performed in isolated adult WT CMs also led to reactivation of the cell cycle and proliferation but only following NRG-1 stimulation (***Figure 2F and G***), as observed in the transgenic *efnb1* cKO model. Finally, NRG-1-induced CM proliferation was confirmed *in vivo* upon NRG-1 injection in 2-month-old mice (Supplementary material online, ***Figure S3A***). NRG-1 treatment did not impact cardiac function (***Figure S3B***), morphometry (***Figure S3C***) or fibrosis (***Figure S3D***), but it significantly induced mitosis (*Aurkb*^*+*^*/DAPI*^*+*^) of CMs (*TroponinT*^*+*^*-cTnT*^*+*^) (***Figure S3E***) as well as cytokinesis (*Aurkb*^*+*^*/Cleavage furrows*) (***Figure S3F***) in cKO mice compared to WT. Mitosis of resident CMs was qualitatively confirmed by nuclear co-staining of Aurora-B with pericentriolar material protein PCM1 (Aurkb^+^/PCM1^+^ cells), a specific CM nuclear marker (***Figure S3G***), despite CM proliferation undervaluation using this marker (***Figure S4*)**.

Overall, these results demonstrate that ephrin-B1 naturally blocks the proliferation potential of resident adult CMs.

### Ephrin-B1 deletion in CMs promotes cardiac regeneration following apectomy in adult mice

To determine whether the proliferation potential of adult CMs lacking ephrin-B1 could be mobilized and ultimately contribute to cardiac regeneration *in vivo*, a natural process restricted to lower vertebrates^27^, we challenged *efnb1 c*KO mice using the radical surgical model of apectomy^28^ at non-regenerative adulthood (2-month-old). WT or cKO mice underwent similar surgical and calibrated resection of the left ventricle apex (**See Methods, *Figure 3A***, Supplementary material online, ***Figure S5A and B***). As expected, 21 days after apectomy, analysis of HE- and Trichrome-stained heart cross-sections from WT mice clearly showed a significant fibrotic scar replacing the excised tissue area (***Figure 3B and C***, Supplementary material online, ***Figure S5C***) devoid of CMs (***Figure 3D***), with a disorganized vasculature (***Figure 3E***). Intriguingly, the resected apex in cKO hearts was replaced by almost normal myocardial tissue with the remarkable presence of a large number of CMs compared to WT (***Figure 3B and D***), considerably reduced fibrosis (∼-50 %, ***Figure 3C***) and highly organized microvasculature (***Figure 3E***). Some CMs detected in the resected area of cKO hearts had reentered the cell cycle and undergone cell division, as indicated by the significant increase of CMs (TroponinT^+^ or PCM1^+^ cells) positive for BrdU (*replication*; ***Figure 3F***), nuclear aurora kinase B (*mitosis*; ***Figure 3G and H***) and aurora kinase B in the cleavage furrows (*cytokinesis*; ***Figure 3I***). CM proliferation in cKO mice was not restricted to the resected tissue area. Indeed, a significantly larger number of smaller (Supplementary material online, ***Figure S5D and E***) and BrdU- or aurora kinase B-positive CMs was identified at the border of the resected area but also at distance in both the ventricles and the septum (Supplementary material online, ***Figure S5F* and *G***). These findings suggest that the apex resection promoted a global tissue response, most likely to compensate for the dynamic ventricular systolic twist defect, in which the apex plays a fundamental role^29^. The cardiac tissue regeneration detected in the *efnb1* cKO mice was consistently associated with a lesser cardiac dysfunction as assessed by echocardiographic left ventricular ejection fraction (***Figure 3J***) and global longitudinal strain (***Figure 3K***).

Overall, these results indicated that ephrin-B1 is a natural blocker of adult CM proliferation *in vivo* that prevents adult cardiac tissue regeneration.

### AAV9-based-Ephrin-B1 knockdown in adult mice hearts promotes cardiac regeneration following myocardial infarction

From a better translational therapeutic perspective, we next determined whether *efnb1* deficiency could similarly boost myocardial repair after myocardial infarction (MI). However, we could not use the *efnb1*cKO genetic model since these mice were highly prone to death compared to WT after MI (Supplementary material online, ***Figure S6A***), reflecting a higher susceptibility to cardiac mechanical stress, as we previously reported^23^. To bypass this problem and to preserve CM adhesion which requires Ephrin-B1 (Supplementary material online, ***Figure S7***), we deleted Ephrin-B1 after the MI induction, based on an adeno-associated virus (AAV) serotype 9-delivery strategy with high cardiac tropism and long-term expression. Two-month-old C59/BL6 mice underwent permanent ligation of the left anterior descending coronary artery and, 3 days post-MI, mice exhibiting 30-40 % left ventricular ejection fraction were selected and immediately injected retro-orbitally with an AAV9 vector expressing shmiR*efnb1* or a control vector (AAV9-mCherry) (***Figure 4A, B***). Efficiency of Ephrin-B1 knockdown was successfully validated *in vitro* but also *in vivo* in the heart at the end of the protocol (Supplementary material online, ***Figure S6B, C***). As shown in ***Figure 4C***, *efnb1* knock-down considerably favored immediate and long-term mouse survival up to 56 days after AAV9 infection compared to control. Left ventricular ejection fraction (LVEF) was significantly stabilized 56 days post-MI in infarcted mice upon *efnb1* knock-down (AAV-shmiR*efnb1*) with some animals exhibiting full LVEF restoration compared to control infarcted mice (AAV-mCherry) that functionally decompensated over time (***Figure 4B***, Supplementary material online, ***Figure S6D***). Histological analysis of the whole hearts showed a significant fibrosis reduction (∼40 %) in AAV9-shmiR*efnb1* infected mice (***Figure 4D***), indicative of an infarct scar size reduction. The increase in heart weight-to-tibia length ratio after MI compared to sham was also significantly higher in these mice, suggesting higher hypertrophy compensation (***Figure 4E***). However, 56 days after *efnb1* knockdown, analysis of the left ventricle (LV) remote zone in the MI group indicated the presence of smaller CMs compared to control AAV9 infection (***Figure 4F***) and the presence of mitotic troponin T-(***Figure 4G***) or PCM1- (***Figure 4H***) positive CMs also undergoing cytokinesis (***Figure 4I***), most likely supporting cardiac compensation through resident CM proliferation. It is noteworthy that CM proliferation could also be measured at distance from the injury site in the septum (Supplementary material online, ***Figure S6E, F***). In all cases, LVEF in AAV9-infected mice negatively correlated with fibrosis and CM area (Supplementary material online, ***Figure S6G, H***).

All together, these data indicate that cardiac deletion of Ephrin-B1 immediately after MI is beneficial for cardiac function by promoting resident CM proliferation and thus cardiac regeneration.

### *Efnb1* KO mice compensate for cardiac senescence through resident CM proliferation

Despite we did not detect CM proliferation in young adult 2-month-old *efnb1* cKO mice, we finally investigated the long-term consequences of *efnb1* deletion, by examining the cardiac phenotype of 12-month-old *efnb1* KO mice. In fact, given that we previously reported that the cardiac tissue disorganization was only focal at 2 month of age^23^, we hypothesized that it could generalize over lifetime and could ultimately lead to cardiac contractile function defect, thus offering the opportunity to use the proliferation potential of *efnb1*-KO CMs as a compensatory mechanism.

At the tissue level, as expected, the absence of ephrin-B1 promoted a general disorganization of cardiac tissue architecture, including the loss of the rod-shape of adult CMs (***Figure 5A***) in all heart compartments (Supplementary material online, ***Figure S8A***), with the presence of whorled CMs in some tissue areas (Supplementary material online, ***Figure S8B***), without fibrosis (***Figure 5B***). Both wild-type (WT) and KO aged mice similarly reactivated the fetal gene program indicative of cardiac hypertrophy remodeling (***Figure 5C***), most likely reflecting a compensatory mechanism for aging stress, as suggested by the specific and substantial breakdown of TGFβ3 inflammatory gene expression (***Figure 5D***), a marker of cardiac senescence^30^. Intriguingly, while only a modest septal hypertrophy could be detected in aged WT mice (IVSTD, ***Figure 5E***), KO mice compensated through a more pronounced septum and left ventricle hypertrophy (***Figure 5E***), but without alteration of the contractile function in both genotypes (***Figure 5F***). Accordingly, old KO mice exhibited a mild but significant increase in heart/body weight ratio, compared to age-matched WT mice (***Figure 5G***). At the cellular level, 3D-stereology-based analysis confirmed 2-D heart cross-section analysis (Supplementary material online, ***Figure S9A***) and revealed a significantly higher CM density (***Figure 5H***) and a reduction in CM volume in old KO mice (***Figure 5I***) compared to their WT counterparts, for which the results were consistent with previous reports when considering age and strain genetic background (*see Methods*). Overall, these results demonstrated that adult *efnb1* KO mice compensated for aging stress through atypical CM hyperplasia, thereby evoking proliferation.

**Figure 5.**
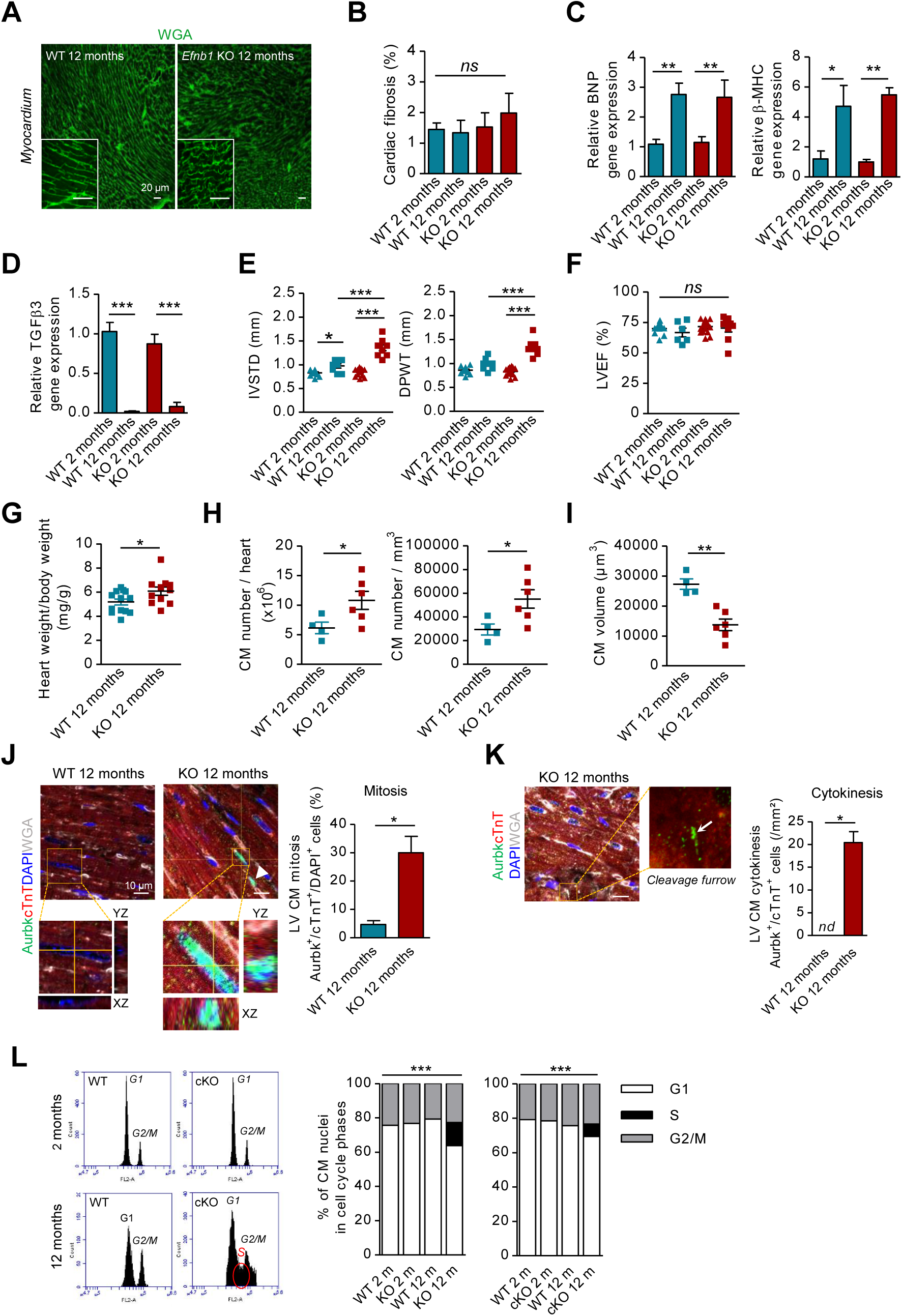
Aged *efnb1* KO mice compensate for aging stress through CM proliferation. **A**, CM morphology within cardiac tissue from 12-month-old WT and *efnb1*^*-/-*^ KO mice stained with fluorescent WGA. **B**, Cardiac fibrosis quantification (n=4 to 6 mice/group). **C, D**, RT-qPCR quantification of BNP, β-MHC (**C**) and TGFβ3 (**D**) gene expression (n=4 to 7 mice/group). **E, F**, Analysis of echocardiography-based morphometry (**E**) and function (**F**) (LVEF, left ventricle ejection fraction) (n=9 to 18 mice/group). **G**, Heart to body weight ratios (n=11 to 13 mice/group). **H, I**, Stereology-based quantification of CM density (**H**) and volume (**I**) (n= 4 to 6 mice/group). **J, K** *In situ* quantification of CM mitosis (**J**, Aurkb^+^/cTnT^+^/DAPI^+^) and cytokinesis (**K**, Aurkb^+^ cleavage furrows/cTnT^+^) (n=3 for each group). **L**, Quantification of cell cycle activity by FACS in propidium iodide-stained nuclei from CMs isolated from 2- or 12-month-old global *efnb1*^*-/-*^ (KO), CM-specific *efnb1*^*-/-*^ KO (cKO) mice and their WT littermates (n=4 mice/group). Data are presented as mean ± SEM and analyzed using Student’s *t*-test for 2 group comparisons or one-way ANOVA with Tukey’s post hoc test for 4 group comparisons * *P*<0.05, ** *P*<0.01, *** *P*<0.001. ns, not significant. nd, not detectable.

The specific capability of adult CMs to proliferate during aging in the absence of ephrin-B1 was confirmed both *in situ* and *in vitro* (see *Methods*). First, in the cardiac tissue, the use of two different specific CM markers (*cardiac troponin T, cTnT; or pericentriolar material protein PCM1*) indicated substantial co-staining with mitosis (*nuclear aurora kinase B-Aurkb*^*+*^*/DAPI*^*+*^) or cytokinesis (*aurora kinase B-Aurkb*^*+*^*/cleavage furrows*^*+*^) markers (***Figure 5J and K***; Supplementary material online, ***Figure S9B***). Second, in isolated CMs (∼ 80 % purity) from aged mice, a notable induction of CM-specific proliferation marker gene expression^26, 31^ from all phases of the cell cycle was found only in KO mice (Supplementary material online, ***Figure S9C***). Finally, flow cytometry analysis of propidium iodide-stained nuclei isolated from purified CMs showed a significant number of aged KO CMs in the DNA replicative S-phase (***Figure 5L***; 13.50±3.05 %), compared to 2- and 12-month-old WT or 2-month-old KO mice that were instead blocked in the G0/G1- and G2/M-phases, as expected due to their post-mitotic state ^25^. These results were confirmed in old CM-specific-KO mice (cKO, ***Figure 5L***; 7.20 ± 0.91 %), thus precluding the contribution of other cell types to this process and clearly validating adult CM proliferation in the absence of ephrin-B1. Noteworthy, DNA replication, mitosis or cytokinesis cell cycle events were not detected in CMs from young 2-month-old KO mice (***Figure 5L***), thus indicating a specific adaptation of *efnb1* KO mice over time, most likely to compensate for aging stress, and not a constitutive CM proliferation phenotype of the *efnb1* KO mice at birth as previously demonstrated.

### Ephrin-B1 sequesters Yap1 at the lateral membrane to prevent adult CM proliferation

We next sought to understand the molecular mechanisms supporting ephrin-B1 as a suppressor of adult CM proliferation. Given its importance in the control of CM proliferation^32^, we examined the Hippo pathway in the 2-month-old *efnb1* cKO mice and more specifically its core effector, Yap1. Yap1 gene (***Figure 6A***) or Yap1 protein expression and activity (phosphorylation state, P-Yap) (***Figure 6B***; *specificity of Yap1 immunostaining in western-blot was assessed as shown in* Supplementary material online, ***Figure S10***) were similar in CMs from resting young cKO or WT mice, with predominance of inactive P-Yap1, thus corroborating the absence of constitutive CM proliferation in *efnb1* cKO mice at resting state. In tissue from WT mice, inactive P-Yap1 was expressed not only at the CM intercalated disk but also at the lateral membrane (***Figure 6C***), where it colocalized with ephrin-B1 (***Figure 6D***). In contrast, in cKO mice, while still expressed in the intercalated disk, P-Yap1 was absent at the lateral membrane, and heterogeneous cytosolic P-Yap1 staining could be observed (***Figure 6C***, transversal view). Given that neither Yap1 global expression nor its activation state were modified in CMs from cKO mice, these results indicated that P-Yap1 was delocalized from the lateral membrane to the cytosol in the absence of ephrin-B1. Noticeably, active Yap1 was absent from CM nuclei (Supplementary material online, ***Figure S11***), in agreement with a lack of CM proliferation in resting cKO mice. The effect of *efnb1* deletion on P-Yap1 delocalization is direct, since P-Yap1 coimmunoprecipitated with ephrin-B1 in WT CMs (***Figure 6E***). The propensity of Yap1 to interact with ephrin-B1 was further confirmed by BRET-based experiments measuring direct protein-protein interactions in real time, in living HEK293T cells (*see Methods***; *Figure 6F***). Altogether, our results showed that CMs from *efnb1* cKO mice harbored a new cytosolic pool of delocalized inactive P-Yap1 that might be mobilized and thus offer a potential for proliferation under stress through Yap1 activation and nuclear translocation. We validated this hypothesis both *in vitro* and *in vivo* since NRG-1, aging and apectomy stimuli significantly promoted Yap1 nuclear translocation (***Figure 6G-I***), as well as the expression of the Yap1-specific proliferation gene target^33^ (***Figure 6J***), only in CMs from cKO. Taken as a whole, these data indicate that ephrin-B1 sequesters inactive P-Yap1 at the lateral membrane of adult CMs, thereby preventing stimulus-promoted Yap-dependent proliferation of CMs.

**Figure 6.**
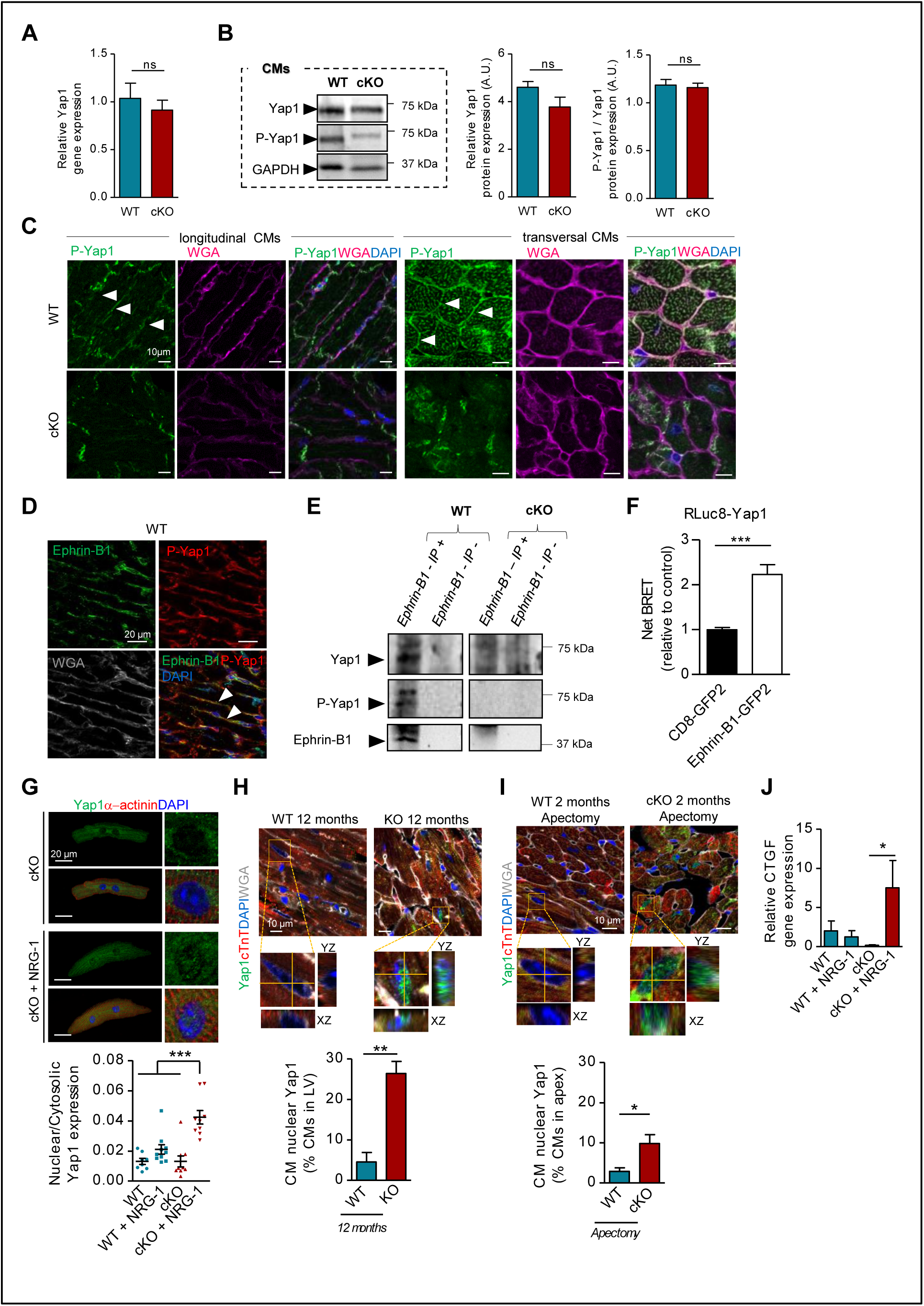
Ephrin-B1 sequestrates P-Yap1 to prevent adult CM proliferation. **A**, RT-qPCR quantification of Yap1 gene expression in isolated CMs from 2-month-old mice (n=4 mice/group). **B**, Western-blot quantification of Yap1 and P-Yap1 (S127) relative to GAPDH expression in CMs isolated from 2-month-old mice (WT n=6; cKO n=5). **C**, P-Yap1 localization at the CM lateral membrane (WT-arrow heads) and lost in cKO. **D**, P-Yap1 and ephrin-B1 colocalization at the CM lateral membrane (arrow heads) visualized on heart cryosections (myocardium) from 2-month-old mice. **E**, Coimmunoprecipitation (IP) assay of Yap1 and P-Yap1 with ephrin-B1 in isolated CM lysates from 2-month-old mice (*IP+: with anti-ephrin-B1 antibody; IP-: without anti-ephrin-B1 antibody*) (n= 3). **F**, Direct interaction between ephrin-B1 and Yap1 measured by BRET in HEK-293T cells (negative control: transmembrane CD8 protein) (Yap1/ephrin-B1 n= 9; Yap1/CD8 n= 4). **G**, Quantification of Yap1 activation evaluated by immunofluorescence of nuclear over cytosolic Yap1 ratios in CMs isolated from 2-month-old mice following or not 8 days of NRG-1 treatment (WT n=8; WT+NRG-1 n=10; cKO n=9; cKO+NRG-1 n=9 cells). **H, I**, Quantification of Yap1 activation evaluated by immunofluorescence of CM nuclear Yap1 (Yap1^+^/cTnT^+^/DAPI^+^) in paraffin sections from **(H)** 12-month-old (WT n=4; KO n=3) or **(I)** 2-month-old apectomized (WT n=4; cKO n=3) mice. **J**, Quantification of Yap-1 target gene expression (CTGF) in isolated CMs from 2-month-old mice following or not 8 days of NRG-1 treatment (WT n=4; WT+NRG-1 n=3; cKO n=5; cKO+NRG-1 n=6). Data are presented as mean ± SEM and analyzed using Student’s *t*-test for 2 group comparisons or one-way ANOVA with Bonferroni post-hoc test for 4 group comparisons * *P*<0.05, ** *P*<0.01, *** *P*<0.001. ns, not significant.

Thus, our observations uncovered a new link between determinants of the adult CM morphology and CM proliferation.

## Discussion

Despite early proliferation arrest of the CM during postnatal maturation coinciding with the establishment of its typical adult rod-shaped morphology, whether these two processes are interconnected is totally unknown. Here, we demonstrated for the first time an essential and dual role of ephrin-B1 in coordinating the morphology and proliferation of the adult CMs by a mechanism relying on a ephrin-B1 scaffolding inactive P-Yap1 at the lateral membrane and thereby controlling the post-mitotic state of this cell (***Figure 7***). In the absence of ephrin-B1, inactive P-Yap1 is released into the cytosol. Under cardiac stress stimuli (apectomy, MI, aging), cytosolic P-Yap1 can then be activated, leading to its translocation into the nucleus to promote transcription of proliferation genes. Ultimately, this signaling cascade leads to striking cardiac compensation through CM proliferation and heart regeneration.

**Figure 7.**
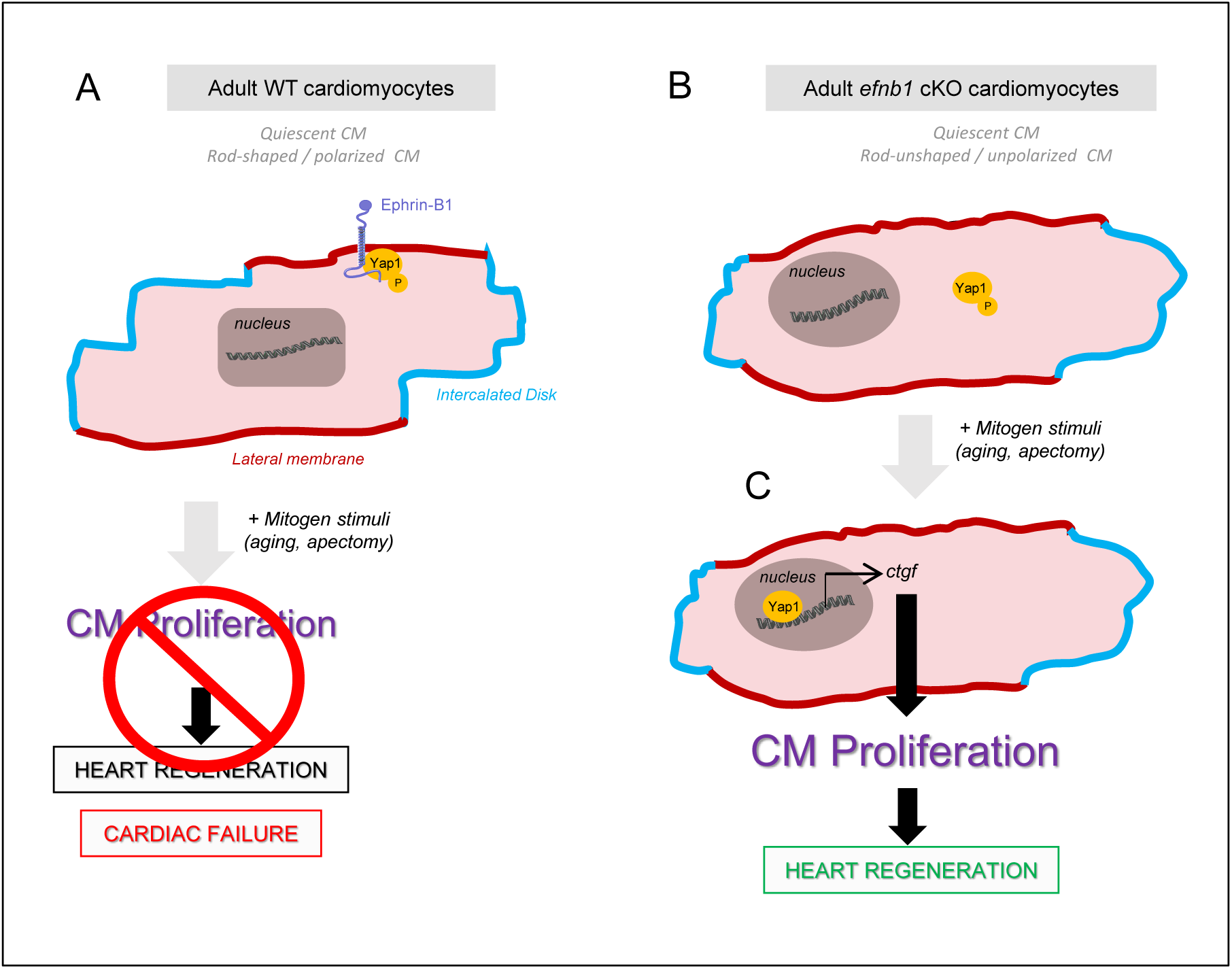
A model for ephrin-B1 coordinating the adult CM morphology and Hippo-dependent proliferation blockage at the lateral membrane. **A**, At the adult stage, ephrin-B1 is complexed at the CM lateral membrane with inactive phospho-Yap1 (P-Yap1), thus preventing potential Yap1-dependent CM proliferation. **B**, Loss of ephrin-B1 in the adult CM releases P-Yap1 in the cytosol, leading to an unpolarized (loss of the rod-shape) but quiescent CM^23^. **C**, However, upon mitogenic cues (still unknown) generated in response to cardiac stress (apectomy, MI, ageing), cytosolic P-Yap-1 can be now mobilized and activated, thus leading to its translocation into the nucleus to promote the transcription of the Yap-1 target *ctgf*-proliferation gene and the ensuing adult CM proliferation and cardiac tissue regeneration.

A major finding of our study was the lack of CM proliferation at resting state in the absence of ephrin-B1 while a substantial proliferation could be triggered to compensate for various cardiac stresses (apectomy, MI, senescence). This observation most likely supports a role for the ephrin-B1-Yap1 pathway as a natural sarcolemmal fine tuner of adult CM proliferation. Consistent with this assumption, re-expression in adult CMs of cell cycle cyclin genes that are naturally downregulated during the CM postnatal maturation was recently shown to lead to up to 15-20 % of CM division following myocardial infarction^34^. Interestingly, CM cytokinesis measured in the aged *efnb1* KO mice also correlated with cyclin re-expression in these cells (Supplementary material online, ***Figure S8C***), in agreement with a vigorous revival of cell cycle progression and high CM proliferation rates in this model. Thus, ephrin-B1 could act as a natural blocking agent of cyclins during the postnatal maturation since ephrin-B1-Yap1 complex expression at the CM lateral membrane coincides with cyclin shutdown and CM proliferation arrest.

In these last few years, the Hippo pathway has been identified as a master regulator of CM proliferation and inactivating the Hippo pathway or activating its effector Yap1 at the adult stage substantially reactivated CM proliferation and cardiac repair after injury^6, 13, 35, 36^. Despite the molecular mechanisms governing the Hippo signaling specifically in the adult CM remaining poorly understood, compartmentalization of Yap1 has emerged recently. Thus, adherent CM junction Fat4 was shown to spatially sequester amotl1-Yap1 and to inhibit Yap activity, thus preventing CM proliferation at neonatal stage, but its role in the adult stage remains unknown^37^. Specific ablation of αE/αT-catenin in adult CMs had no impact on the CM proliferation state in unstressed hearts but favored cardiac regeneration after myocardial infarction through CM proliferation and Yap nuclearization^14^, emphasizing a fundamental role of the intercalated disk for Yap availability in the adult CM, but one that is mechanistically unknown. More recently, dystroglycan 1, a component of the DGC complex, localized at the lateral membrane of adult CMs and connecting the ECM and the contractile apparatus, was identified as a new, direct partner and repressor of Yap activity since mice with inactive DGC (*Mdx* mice) constitutively promoted CM proliferation through a Yap-dependent pathway^38^. In contrast, we demonstrated here new, original regulation of the Hippo pathway at the CM lateral membrane, relying on the spatial sequestration of inactive P-Yap1 by ephrin-B1 with no impact on its activity. This result provides a new level of information to understand the incompetence of the adult CM to proliferate relative to the Hippo pathway since, up to now, it has been related to the activation of the Hippo pathway and inactivation of its effector, Yap1. Here, we demonstrated that the single spatial sequestration of inactive P-Yap1 at the lateral membrane could prevent its activation for further downstream proliferative signaling events. These results revealed the coexistence of multiple Hippo pathways at the lateral membrane of the adult CM. Consistently, we previously characterized ephrin-B1 as a new protein of the lateral membrane acting independently from the DGC or integrin systems but also from the contractile apparatus^23^. In these last few years, similar Yap subcellular sequestration has emerged as a new, original mechanism to regulate the Hippo pathway in other models^39^. Moreover, different Yap-dependent signaling pathways have been shown to coexist in the same cell^40^, highlighting the complexity of this system. Similar complexity does apply to the adult CM.

How can we now reconcile ephrin-B1 and Yap1 interaction from a structural standpoint? Ephrin-B1 can signal through its intracellular domain, which can recruit various signaling proteins through both PDZ- and tyrosine phosphorylation-dependent interactions^41^, thus precluding any direct interaction with Yap1 regulatory domains^42^. However, we previously identified an atypical complex at the lateral membrane of the adult CM formed between Ephrin-B1 and two tight junctional components, claudin-5 and Zo-1^23^. Given that Yap-1 can directly interact with Zo-1 through its PDZ binding domain, ephrin-B1 must be part of a larger complex that acts as a Yap-1 scaffolder.

Finally, we provide a new molecular mechanism with ephrin-B1 acting as a hub connecting cell morphology and proliferation. Interestingly, zebrafish retains cardiac regenerative properties through proliferation of resident CMs^43^ and their ventricular CMs harbor a pseudo-rod shape (spindle-shaped)^44^, thus reminiscing ephrin-B1 defective adult-CMs but also abortive differentiated mammalian CMs as indicating by the lack of T-Tubules, a marker of lateral membrane invaginations. Whether these CMs do not express ephrin-B1 homolog is completely unknown but could be an attractive hypothesis connecting immature rod-shape CM to their proliferative benefit. Consistent with this hypothesis, round-shaped neonatal CMs proliferate and do not express ephrin-B1 predominantly at their plasma membrane^23^.

Overall, our study provided the first demonstration of the existence of a molecular scaffold at the lateral membrane of the adult CM, *i*.*e*. ephrin-B1, connecting the cell cycle arrest and morphological features of this cell. Finally, this study demonstrates that therapeutic targeting of ephrin-B1 in the adult heart could represent an attractive new strategy in regenerative cardiac medicine using resident CMs for the treatment of post-ischemic patients.

## Acknowledgements

We thank S. Meilhac (Imagine-Pasteur group, Paris, France) for critical reading of the manuscript and helpful discussions and A. Davy (Centre de Biologie du Développement, CNRS, Toulouse, France) for kindly providing us with the global *efnb1* KO mice. We thank the ANEXPLO core facility for assistance with echocardiography, the TRI Genotoul Network core facilities for assistance with imaging (M. Vigneau for the stereology study), and the I2MC histology (L. Fontaine, UMR 1048 INSERM) core facility. We are also grateful to S. Estaqué and N. Loukh (Département d’Histopathologie, Toulouse University Hospital) for the 60 µm thick paraffin-embedded heart sections for the stereology study.

## Supplementary material

### Supplementary Methods

#### Isolation and culture of adult CMs

Isolation of adult CMs from mice was performed using the Langendorff perfusion method as previously described (1). After isolation, dissociated CMs were plated on laminin 2 (10 µg/ml)-coated culture dishes. After 15 min, plating medium was changed to culture medium containing MEM, 1 % penicillin-streptomycin-glutamine, 1 % insulin-transferrin-selenium, 4 mM NaHCO_3_, 10 mM Hepes, 0.2 % BSA and 25 μM blebbistatin which was refreshed every day.

#### *In vitro* adult CM proliferation assay

Isolated adult CMs were cultured in their cell culture medium in the presence or not of 100 ng/ml neuregulin-1 (NRG-1, EGF-like domain, amino acids 176–246; R&D Systems, 396-HB-050) for 8 days. The NRG-1-containing medium was refreshed every day. For detection of DNA synthesis, 5-bromo-2-deoxyuridine labeling reagent (BrdU, 1:100 dilution, Life technologies, 00-0103) was supplemented to the culture medium every day from the second day of CM culture.

#### CM nucleation

Immediately after adhesion to the laminin 2-coated plates, isolated ventricular CMs were fixed in 4 % paraformaldehyde (PFA) and co-stained with 4’,6’-diamidino-2-phenylindole (DAPI, Sigma Aldrich, 32670) and Oregon green® 488 conjugated Wheat-Germ-Agglutinin (WGA, Life technologies, W6748). Images were acquired on a Zeiss Observer Z.1 microscope and the nucleation profile was manually quantified using Axio Vision Rel 4.8 software.

#### Immunocytochemistry

CMs were fixed in ice-cold methanol fixation, permeabilized (0.1 % Triton X-100) and blocked in 1 % bovine serum albumin, CMs were immunostained using standard protocols with the following primary antibodies (all diluted in 0.1 % BSA / 0.1 % Tween 20 / 0.5 % Triton X-100 PBS) incubated overnight at 4°C: mouse anti-α-actinin (Sigma Aldrich, A7732), rat anti-BrdU (Abcam, ab6326), rabbit anti-aurora kinase B (Abcam, ab2254), rabbit anti-histone H3 phosphorylated at serine 10 (pH3, Cell signaling, 9701S), rabbit anti-acetyl-histone H3 (H3K9/14Ac, Millipore, 06-599), rabbit anti-tri-methyl-K9 histone H3 (H3K9me3, Abcam, ab8898), goat anti-Ephrin B1 (R&D, AF473), rabbit anti-Yap1 (Cell signaling, 14074). Cells were washed and then incubated with the corresponding secondary antibodies conjugated to specific fluorophores (Alexa-Fluor 488 goat anti-rabbit Ig-G, Alexa-fluor 594 donkey anti-rabbit IgG, Texas-red goat anti-mouse IgG or Alexa-fluor 488 donkey anti-rat IgG, Life technologies). Nuclei were visualized with DAPI (Sigma Aldrich, 32670). Fluorescent images were acquired on a Zeiss Observer Z.1 microscope with *Apotome* optical sectioning using Axio Vision Rel 4.8 software (Carl Zeiss). For Yap1 expression studies in isolated adult CMs, images were acquired in XYZ on a Zeiss LSM 780 confocal microscope using Zen 2011 software (Carl Zeiss).

#### Quantification of Yap1 activation *in vitro*

Yap1 activation (nuclear translocation) was measured as the ratio of nuclear over the cytosolic Yap1 immunofluorescence pool. Nuclear to cytosolic Yap1 immunofluorescence ratio in purified CMs in culture was conducted using a Fiji home-made macro relying on both Yap1 and DAPI immunofluorescence detected in different z-stacks / cell. For each cell, only z-stacks in which nuclei were visible have been quantified individually. The results represent the ratio between quantification of nuclear Yap1 (sum of all quantified z-stacks) to cytosolic Yap1 (nuclear Yap1 subtracted to total Yap1 calculated from the sum of all quantified z-stacks). Quantifications have been selectively performed on binucleated CMs (major CM population) only to avoid bias.

#### Histology

Hearts were excised and immediately fixed in 10 % formalin (24 h), 4 % formalin (48 h) and then embedded in paraffin and sectioned at 6 μm intervals. Except for aging studies (transversal sections), all other heart sections were longitudinal.

#### Immunohistochemistry

Formalin-fixed paraffin-embedded or frozen tissues were used for immunohistochemical analysis. For paraffin-embedded sections, samples were deparaffinized, rehydrated and subjected or not to heat-induced epitope retrieval depending on the antibodies. For cryosections, samples were fixed with acetone or 4 % PFA. Heart sections were then permeabilized (0.5 % Triton X100-PBS), blocked (Dako Protein Block, Dako, X0909 or 10 % normal goat serum, Dako, X0907 or 0.5% BSA/PBS), and stained overnight at 4°C with the following primary antibodies all diluted in 0.1 % BSA / 0.1 % Tween 20 / 0.5 % Triton X-100-PBS: rabbit anti-troponin I (Santa Cruz biotechnology, sc-15368), mouse anti-troponin T (ThermoFischer scientific, MA5-12960), rat anti-BrdU (Abcam, ab6226), rabbit anti-aurora B kinase (Abcam, ab2254), rabbit anti-Yap1 (Cell signaling biotechnology, 4912), rabbit anti-PCM1 (Santa Cruz biotechnology, 50-164), goat anti-Ephrin-B1 (R&D, AF473), rabbit anti-phosphoYap1 (S127) (Cell signaling Technology, 4911). Secondary fluorescent antibodies used in this study were as follows: Alexa-Fluor 488 goat anti-rabbit IgG, Texas-red goat anti-rabbit IgG, Texas-red goat anti-mouse IgG, Alexa-Fluor 488 goat anti-mouse IgG, Alexa-fluor 488 donkey anti-rat IgG, Alexa-fluor 594 donkey anti-rabbit IgG, Alexa-Fluor 488 donkey anti-rabbit IgG (Life technologies). Nuclei were visualized with DAPI (Sigma Aldrich, 32670). Cell membranes were stained with Texas Red- or Alexa-fluor 633-conjugated Wheat Germ Agglutinin (WGA, Life technologies, W21404). Endothelial cells were stained with OG488-conjugated Griffonia simplicifolia isolectin-B4 (Life technologies, I21411). All Images were acquired on Zeiss LSM 780 confocal microscope using Zen 2011 Software (Carl Zeiss). For Ephrin-B1/P-Yap1, colocalization images were acquired with FRAME mode.

#### Quantification of CM proliferation *in situ*

All quantifications have been conducted in paraffin-embedded sections by confocal imaging as described above in “Immunohistochemistry” and were performed in z to ascertain the labelling in nuclei form resident CMs (Troponin T or I TnT/I-positive cells). To ascertain proliferation of resident CM, we also performed quantifications using pericentriolar material protein PCM1, a specific CM nuclear marker (2), despite this marker underestimates these cells as shown in *SI Appendix*, Fig. S3. In most cases, we also included an additional fluorescent WGA label to mark cell borders for better appreciation of CMs. Both mitosis and cytokinesis events were visualized by the use of aurora kinase B-nuclear labeling (Aurbk+/TnT+/DAPI+ cells) or aurora kinase B labeling on cleavage furrows (non nuclear: Aurbk+/TnT+ cells) respectively since this marker is involved in both cell cycle phases (3). Specificity of anti-aurora kinase B antibody batch was evaluated based on the use of neonatal cardiac tissue (P3) with known proliferation as positive control. Finally, all quantifications were performed by two observers blinded to the sample identity and gave similar results.

#### Quantification of cardiac fibrosis

Fibrosis was quantified on Masson’s trichrome-stained paraffin-embedded heart sections. For apectomy, the apical cardiac fibrosis (3-4 sections at 200 µm intervals/heart) was quantified using NIS Element Basic Research version 2.31 (Nikon imaging software) in the all apex area (between the above subventricle and bottom apex). Similar analysis was conducted for quantification of myocardial fibrosis in the aging mouse models. The experimenter was blinded to the mouse genotype.

#### 2D quantification of CM area/density

For *in situ* quantification of CM surface area and density, deparaffinized heart slides were stained with Texas Red-conjugated-WGA (Life technologies, W21405) and CM area and density were measured in transversal (aged mice) or longitudinal (apectomy) heart cross-sections by manually tracing the cell contour on images of whole hearts acquired on a digital slide scanner *NanoZoomer* (Hamamastu) using Zen 2011 software (Carl Zeiss). The experimenter was blinded to groups and genotypes.

#### Stereology-3D quantification of CM volume/density

Stereology analysis was conducted on 50-60 µm-thick paraffin-embedded heart sections. After deparaffinization and rehydratation, sections were permeabilized with proteinase K (S3020, 3 minutes, room temperature), blocked (Protein Block, Dako, X0909) and incubed overnight at 4°C with Oregon green® 488 conjugated Wheat-Germ-Agglutinin (WGA, Life technologies, W6748) to visualize all cell borders. Z-stack images were then acquired on a Leica TCS SP8 confocal microscope (63x oil immersion objective) up to 40-60 µm with an optimal step of 0.3 µm applying compensation of intensity loss during the z-stack acquisitions. Stereological analysis was performed using Image J software on z-stacks and brightness inversion of the AF488-WGA staining to provide clearer delimitation of CM surface. The number of CMs was counted manually (point function on Image J) on each acquired image (∼ 150 z-stacks). CM density was estimated as the global acquisition volume relative to the CM number. To estimate the CM volume, we proceeded first to image binarization and manual cleaning so to exclude non CM cells. The average CM volume was then estimated as the global CM volume within the image relative to the CM number. The total CM number in the heart was estimated as previously described (4) by using N_v_ × V_REF_ method (N_V_ is the estimated CM density and V_REF_ is the reference volume of the heart). The reference volume was calculated using the tissue density of the myocardium (1.06 g/cm^3^) (4). It should be noticed that calculation of the total CM number / heart takes into account the heart weight which can highly differ depending on the genetic background of the mouse strains(5) but also on animal housing and diet (6). In agreement, 2-month old S129/S4 × C57BL/6 global *efnb1* KO mice used in this study harbor around 150-180 mg heart weights while they are around 90-100 mg range in the C57Bl/6 mice typically used for instance by Alkass et al (7) consistent with 2 fold less CM/heart reported in their stereology study.

#### Analysis of cardiomyocyte cell cycle

Cell cycle was evaluated on purified CM nuclei. Thus, isolated ventricular CMs were first fixed in ice-cold 70 % ethanol and intact nuclei were obtained from 1 mg/ml pepsin / 0.2 M HCl digestion (15 min, 37°C). After purification, nuclei were stained with 200 μg / mL propidium iodide (Life technologies, P1304MP) / 12 μg / mL RNase A (Roche)-PBS and further proceeded for FACS experiments (BD Accuri^™^ C6 flow cytometer and BD Accuri C6 software).

#### Total RNA isolation and real-time quantitative RT-PCR

Total RNA extraction from isolated CMs was performed using Trizol according to the manufacturer’s instructions (TRI Reagent®, Molecular Research Center). RNA quality and quantification of extracted RNA were assessed by an Experion automated electrophoresis system (Biorad). First-strand cDNA was synthesized using the superscript II RT-PCR system (Invitrogen) with random hexamers. Negative controls without reverse transcriptase were conducted to control the absence of genomic DNA contamination. Fifteen nanograms of cDNA from RT reaction were then mixed with specific primers (listed in Supplementary Table 1) and EVA green mix (Euromedex). Real-time PCR was performed in 96-well plates using an ABI 7900 Fast (Applied Biosystems). Geometric mean values of GAPDH and HPRT housekeeping genes were used for normalization, as previously described (1). Melting curve analysis was performed to ensure a single PCR product and a specific amplification. Relative gene expression was calculated using the 2^−ΔΔCT^ method (1).

#### Lentivector constructs, transduction and evaluation of gene down-regulation

Three shRNA constructs targeting Ephrin-B1 (sh*efnb1*) were generated by Sigma-Aldrich with the pLKO.1-puro vector. The 2 (shB1-44, -45 and -47, respectively). The sh*efnb1* and the control shRNA vectors (targeting luciferase, sh*luc*, SHC007, Sigma-Aldrich) were produced using the tri-transfection procedure with the plasmids pLvPack and pLvVSVg (Sigma-Aldrich, Saint-Quentin Fallavier, France), in HEK-293FT cells. They were evaluated for their ability to transduce the C2C12 myoblast cell lines (ATCC, Catalog No. CRL-1775^™^) and adult CM I cultures, both naturally expressing Ephrin-B1. The efficiency of cell transduction *in vitro* was checked in parallel by FACS analysis using the pTRIP-GFP lentivector (8). The efficiency was at least 40 % for C2C12 cells and 99 % for adult CMs. C2C12 cells (4 × 10^4^/well) were plated into 6-well plate and transduced overnight in Dulbecco’s Modified Eagle’s Medium (DMEM) with 20 % fetal calf serum at 37°C with 5 % CO_2_ in the presence of purified concentrated lentiviral vector. Cells were transduced with 5.10^5^ transduction unit (TU) of the three sh*enfb1* lentivectors per well or with 5.10^5^TU of the sh*luc* lentivector as control. Primary CMs were transduced using the same protocol in their specific culture medium. The efficiency of the sh*efnb1* was evaluated by real-time quantitative RT-PCR (specific primers are listed in *SI Appendix*, Table S1) and promoted ∼80 % *efnb1* gene downregulation (not shown). For that purpose, total cellular RNA was extracted from C2C12 cells (endogenously expressing eprhin-B1) and purified using the Trizol reagent (Invitrogen) according to the manufacturer’s instructions. A total of 1 µg RNA was used to synthesize cDNA using a High-Capacity cDNA Reverse Transcription Kit (Applied Biosystems, Villebon s/ Yvette, France). Ephrin-B1 gene expression was investigated using SsoFast Eva Green Supermix (Bio-Rad) and performed on a StepOne sequence detection system (Applied Biosystems). Each reaction was run with HPRT as a reference gene and all data were normalized based on HPRT expression levels.

The Yap1-expressing lentivector was obtained by subcloning the Mus musculus variant 1 Yap1 (Clone sequence: BC041733.1) in the pTRIP lentivector.

#### ShmiR construction and rAAV production

In the knockdown experiments *in vivo*, we used shmiRs as they have been reported to produce siRNAs with more efficiency than shRNAs^10-12^. Indeed shmiRs undergo a better processing and less toxicity than shRNAs. Design and construction of the shmiR targeting Ephrin-B1 was adapted from the BLOCK-iT^™^ PolII Mir RNAi expression system (Stratagene, Massy, France): two complementary oligonucleotides mEB1-47 Top and mEB1-47 Bottom (Supplemental Table 2) containing the sequence targeting mouse Ephrin-B1 with flanking regions derived from miR-155, were annealed, phosphorylated and cloned between the XbaI and BglII sites of the plasmid pAAV-MCS (Stratagene). The resulting recombinant AAV vector, rAAV-shmiR*efnb1* and the control rAAV-*mCherry* (pAAV-MCS, Stratagene) were produced as previously described^9^ by calcium phosphate tri-transfection procedure of HEK293 cells (pAAV-shmiR*efnb1* or pAAV-mCherry and pAAV2/9 and pHelper (Stratagene)). The titers for the recombinant AAV9-*mCherry* and AAV9-shmiR*efnb1* were 9.10^12^ vg/ml and 1.10^13^ vg/ml respectively.

#### MicroRNA isolation and quantification

For miR-195 quantification, RT was performed on 500 ng of total RNA (Trizol extraction) using the miScript (Qiagen) kit. For the real-time RT-PCR reaction, the resultant cDNA was diluted 1:100. Each RTstep was performed in duplicate and the qPCR in triplicate for each reaction. U6 RNA was used as an endogenous control (primer sequences listed in Supplemental Table 1) and relative miR-195 (Qiagen) quantity calculated by the 2^−ΔΔCT^ method. MicroRNA expression was quantified by PCR using a Bio-Rad thermal-cycler. miR-195 primer sequences were purchased from Qiagen and used according to the manufacturer’s protocol.

#### Apical resection (Apectomy)

Two-month-old adult mice were anesthetized (intraperitoneal injection with 125 mg/kg ketamine plus 10 mg/kg xylazine for induction and 0.5 % O_2_ / 1 % isoflurane mixture inhalation for anesthesia maintenance), placed on a heating mat and subjected to artificial ventilation through endotracheal cannulation. The animals were positioned on their right side and a thoracotomy was performed on the sixth or the seventh intercostal space. After exposition of the heart apex, apical resection was controlled using a 3/8 circle needle and 8.0 polypropylene thread and performed with microscissors following the curvature of the thread to ensure surgery reproducibility (*SI Appendix*, Fig. S6*A* and Video S1). Apex resection was normalized by apex weighing (*SI Appendix*, Fig. S6*B*). After hemostasis control, the intercostal space was closed in two separate points using a 6.0 nylon monofilament, Ethnor), followed by skin surface suture. Then, isoflurane was stopped and subcutaneous buprenorphine (100 µg/kg) was administered. Mice remained air ventilated until the return of peripheral reflexes. The whole procedure was performed using an operating microscope (Zeiss OPMI 1 FC). Sham animals underwent exactly the same surgical procedure without apex resection. One day after surgery, mice that had undergone apectomy were indistinguishable from sham-operated ones. Hearts were collected 21 days following apectomy. For quantification of CM replication in cardiac tissue, BrdU was administered by intraperitoneal injection on days 1, 7 and 14 following apectomy (0.01 ml BrdU solution/g; Life technologies, 00-103).

#### Echocardiography

For experiments on aged mice, animals were anesthetized by intraperitoneal injection of 10 μg/g etomidate and underwent noninvasive transthoracic echocardiography using a General Electric Vivid 7® (GE Medical System) equipped with a 14 MHz linear probe. Cardiac ventricular dimensions were measured in M-mode images at least 5 times for each animal. Left ventricle ejection fraction (LVEF) was calculated using the Teichholz formula. For apectomy and NRG-1 *in vivo* experiments, animals were anesthetized with a 0.5 % O_2_ / 1 % isoflurane mixture and underwent noninvasive transthoracic echocardiography using a Vevo® 2100 (VisualSonics). LVEF was measured in long-axis B-mode at least three times for each animal. Global longitudinal strain was measured from parasternal long-axis images using speckle-tracking-based imaging to evaluate global cardiac performance. All measurements were obtained by an examiner blinded to the genotype of the animals.

#### Immunoprecipitation and Western-blotting

For immunoprecipitation (IP) experiments, isolated CMs were lysed in CHAPS buffer (140 mM NaCl / 2 mM EDTA / 25 mM TrisBase, pH 7.4 / 1.5% CHAPS) while for CM protein analysis, isolated CMs were lysed in Triton buffer (150 mM NaCl / 1 mM EDTA / 50 mM TrisBase, pH 7.4 / 1% Triton X-100) both supplemented with complete protease and phosphatase inhibitors (Roche). Protein extracts were immunoprecipitated (1 mg) with anti-Ephrin-B1 (R&D, AF473) or anti-Yap1 (Cell signaling, 4912) antibodies, or directly subjected (50 µg) to SDS-PAGE and transferred into nitrocellulose membranes. Proteins were detected with primary antibodies followed by Horse Radish Peroxydase-conjugated secondary antibodies to goat, mouse, or rabbit IgG (GE Healthcare) using enhanced chemiluminescence detection reagent (GE Healthcare). Protein quantification was obtained by densitometric analysis using ImageJ software and was normalized to GAPDH expression and expressed in arbitrary units (A.U.). Due to visualization of multiple bands around expected Yap1 staining, we used positive (H9C2 over-expressing Yap1) or negative control (siRNA) to ascertain specificity of anti-Yap1 antibody.

#### Bioluminescence Resonance Energy Transfer (BRET)

For BRET experiments, mouse Ephrin-B1 cDNA encoding plasmid was a generous gift from A. Davy (Toulouse. France) and was C-terminally fused in frame to GFP2 in pGFP2-N3 (Perkin). Mouse Yap1 cDNA was obtained from Dharmacon (IMAGE:4239820). Yap1 was fused in frame at the N-terminus region to *Renilla* Luciferase (*R*Luc8) in modified vectors derived from pEGFP-C3 (Clontech). All plasmids were fully sequenced. BRET^2^ was measured between *R*Luc8- and GFP2-fusion proteins following transient transfection of appropriate vectors in living HEK293T cells and in real time as previously described (9). To control for specificity of interactions with transmembrane Ephrin-B1, we conducted BRET experiments by using instead an unrelated transmembrane CD8-GFP2 construct as previously described (9), thus recapitulating the secretory pathway distribution of Ephrin-B1 protein to test for protein crowding/collision effect that could lead to false positive BRET signals.

### Supplementary Tables

**Table S1.**
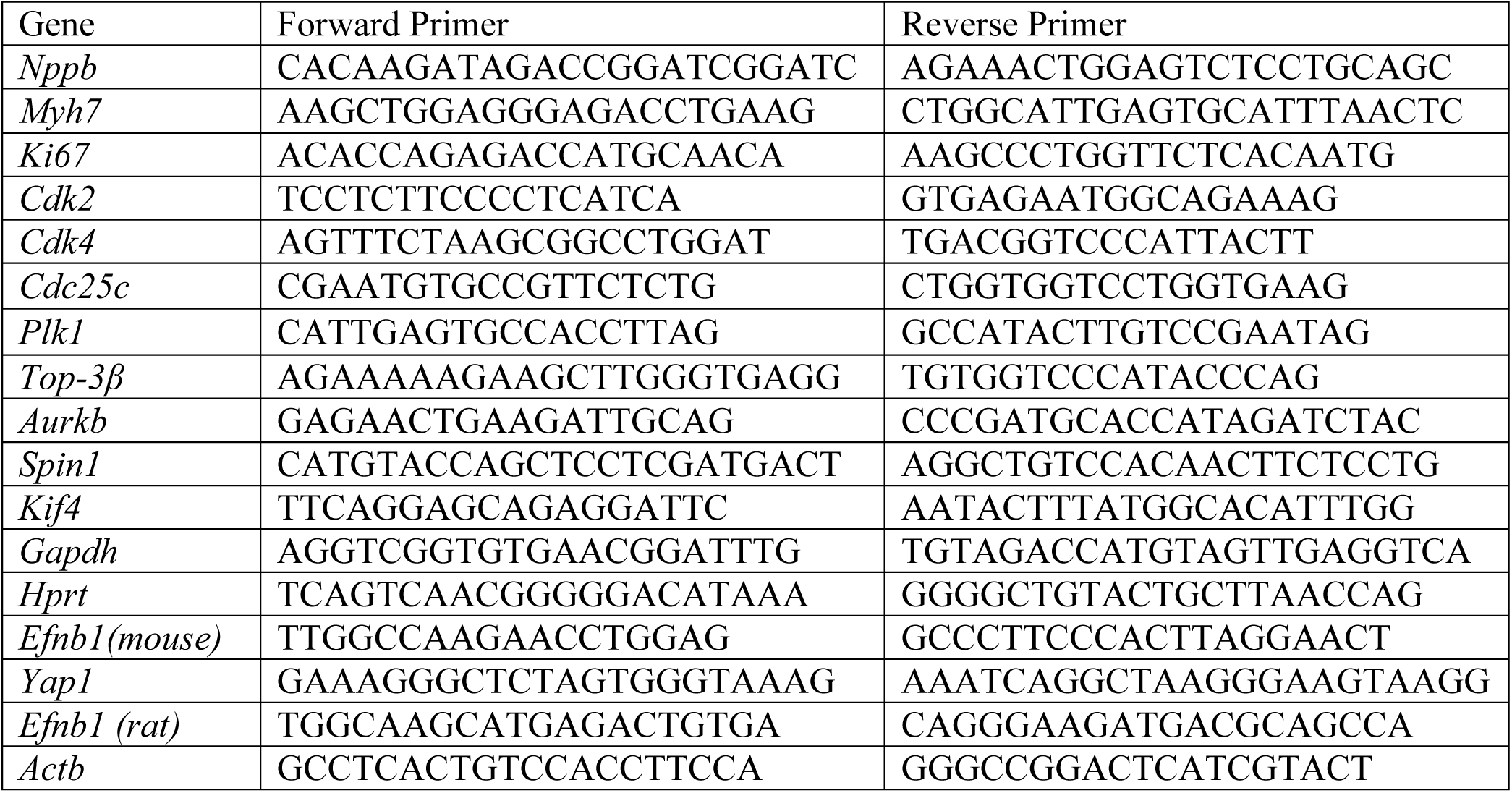
qRT-PCR primer sequences.

**Table S2.**
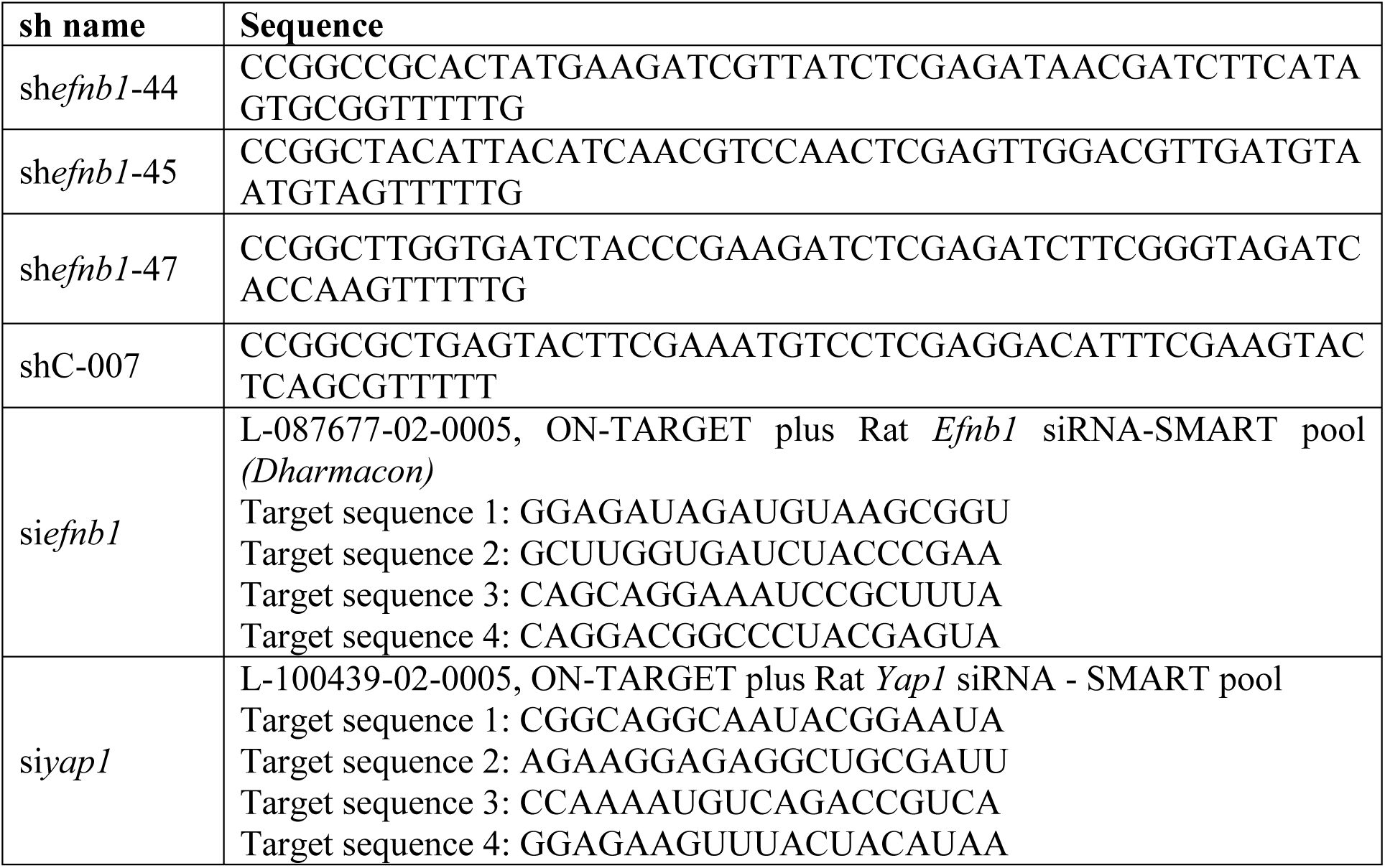
Sh-RNA and si-RNA sequences.

## Supplementary Figure legends and video legend

**Figure S1.**
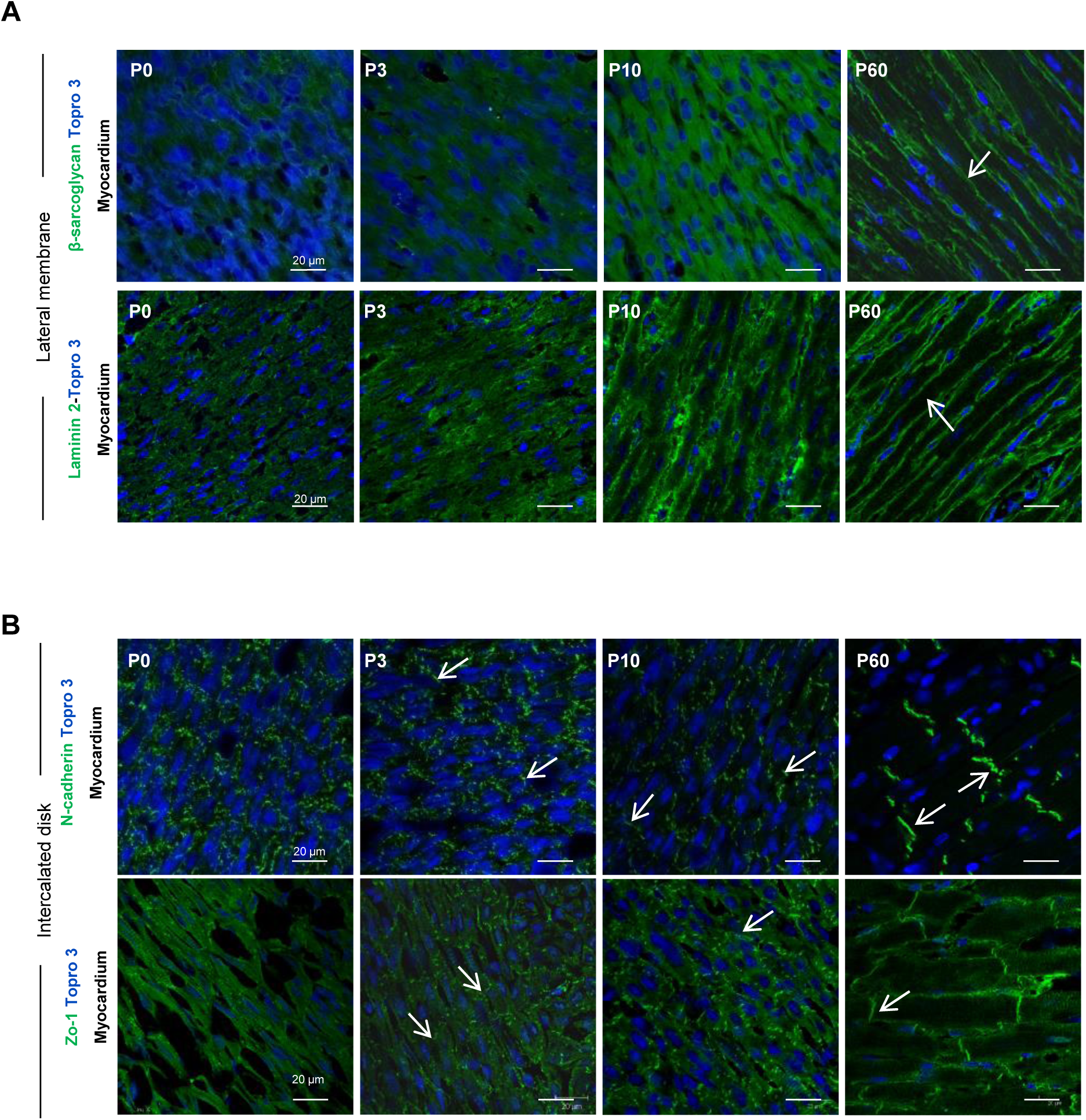
Setting of lateral membrane and intercalated disk determinants during the postnatal maturation of the CM rod-shape. **A-B**, specific markers for (**A**) the CM lateral membrane (β-sarcoglycan and laminin 2) or (**B**), the intercalated disk (N-cadherin and Zo-1) visualized by immunofluorescence on heart cryosections from rats at different cardiac postnatal maturation days (P0, P3, P10, P60). Cell nuclei were immunostained with Topro3 marker.

**Figure S2.**
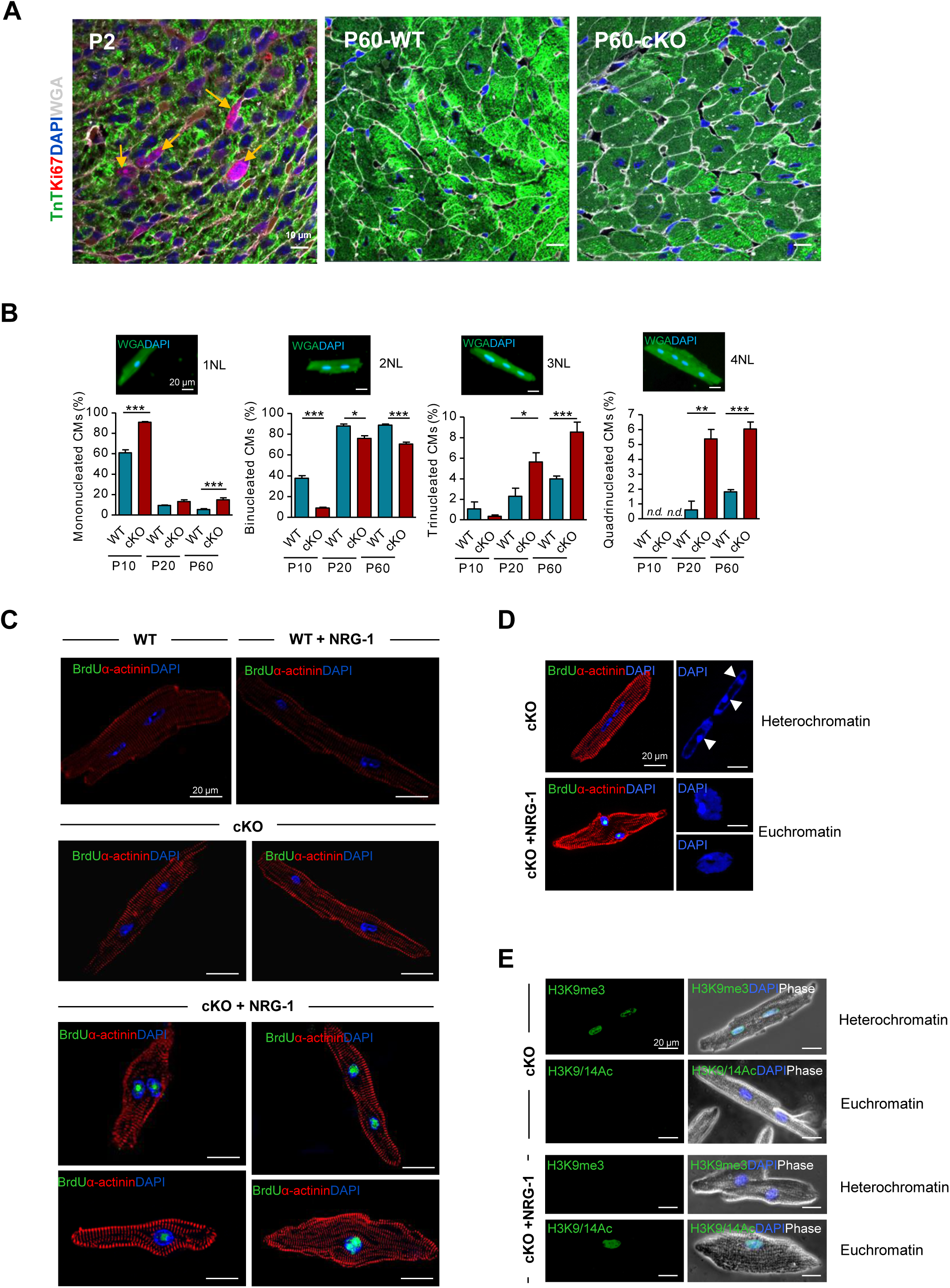
Indirect morphological indicators of CM proliferation under Neuregulin-1 stimulation in the absence of Ephrin-B1. **A**, Evaluation of global cell cycle activity (TnT+/DAPI+/Ki67+) of CMs from the cardiac tissue of postnatal day 2-old mice (positive control) or 2-month-old WT or *efnb1 c*KO mice. **B**, Nucleation profile of CMs isolated from postnatal days 10, 20 and 60 days (P10-P60) (∼350 CMs/mouse, n=4 to 8 mice/group). **C-E**, Representative immunofluorescent imaging of: (**C**) CM replication (BrdU^+^/α-actinin^+^/DAPI^+^) isolated from 2-month-old WT or cKO mice and treated or not with NRG-1 for 8 days. Sarcomere (α-actinin) disarray and nucleus rounding were only observed in NRG-1 treated-CMs from cKO; (**D**) “Active” euchromatin (decondensed) and “inactive” (condensed; arrow heads) heterochromatin evaluated through DAPI- or (**E**) Histone H3 acetylation-(H3K9/14Ac/euchromatin) or methylation-(H3K9me3/heterochromatin) staining of CMs isolated from 2-month-old cKO and treated or not with NRG-1 for 8 days. Euchromatin was only detected in NRG-1-treated CMs. All data are mean ± SEM.; Student’s *t*-test for 2 group comparisons or one-way ANOVA with Tukey post-hoc test for 4 group comparisons; * *P*<0.05, ** *P*<0.01, *** *P*<0.001.

**Figure S3.**
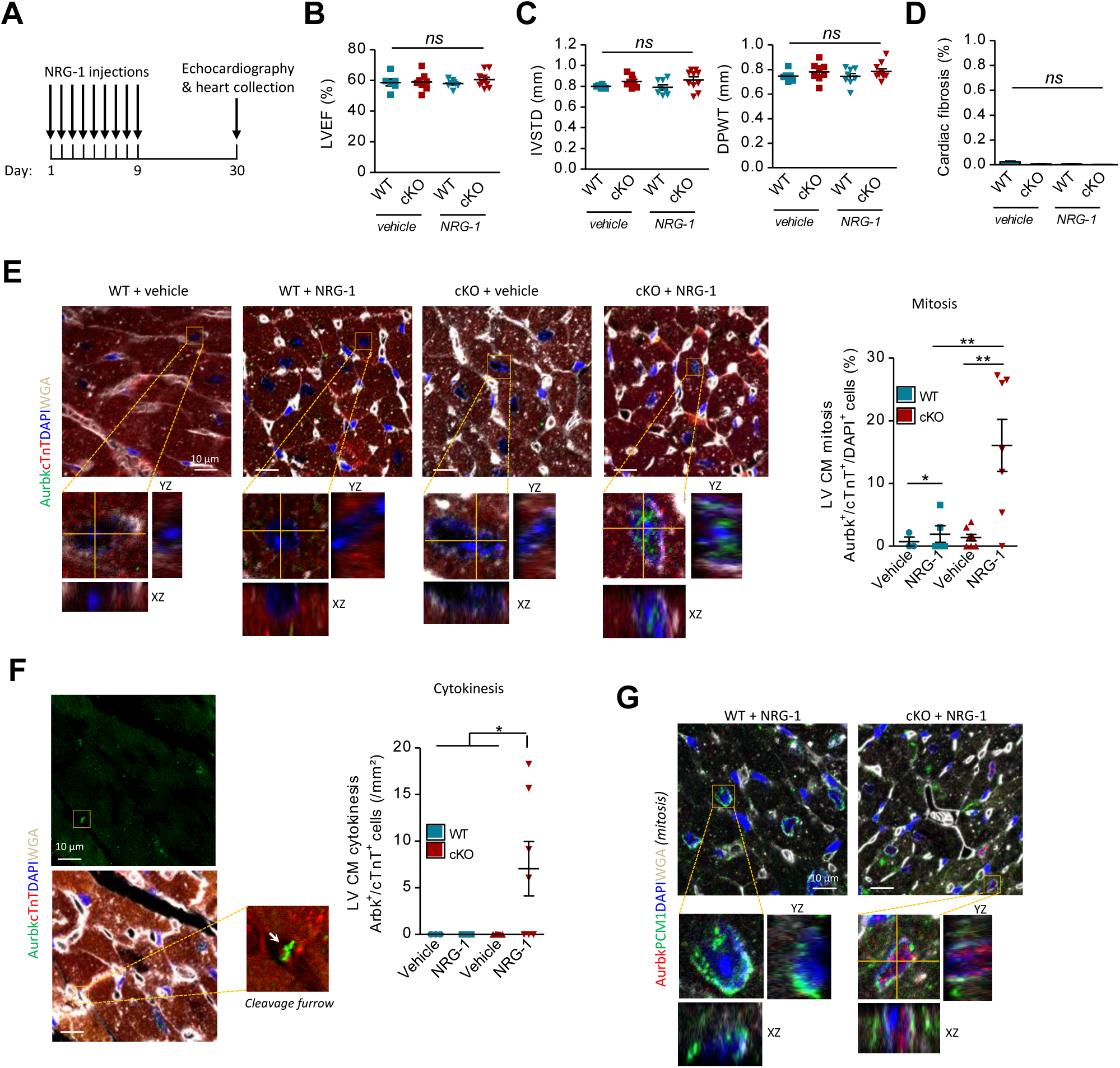
Two-month-old *efnb1 c*KO mice exhibit proliferation-competent cardiomyocytes *in vivo*. **A**, Schematic of the *in vivo* experimental protocol of NRG-1 injections in 2-month-old WT or cKO mice. **B, C**, Analysis of echocardiography-based function (**B**, LVEF, Left Ventricle Ejection fraction) and morphometry (**C, IVSTD**, Interventricular Septal Thickness in diastole, **DPWT** diastolic posterior wall thickness of the left ventricle) (n=6 to 10 mice/group). **D**, Quantification of cardiac fibrosis following 1 month Neuregulin1 (NRG-1) treatment (n=3 mice/group).**E**, *In situ* quantification of CM mitosis (Aurkb^+^/DAPI^+^/cTnT^+^) (n=4 to 7 mice/group). **F**, *In situ* quantification of CM cytokinesis (Aurkb^+^ cleavage furrows/cTnT^+^) (n=4 to 7 mice/group). **G**, Representative immunofluorescent images of resident CM mitosis (PCM1^+^/DAPI^+^/Aurkb^+^) in cardiac tissue from WT or cKO mice treated with Neuregulin1 (NRG1). All data are mean ± SEM.; Student’s *t*-test for 2 group comparisons or one-way ANOVA with Tukey post-hoc test for 4 group comparisons * *P*<0.05, ** *P*<0.01. ns, not significant.

**Figure S4.**
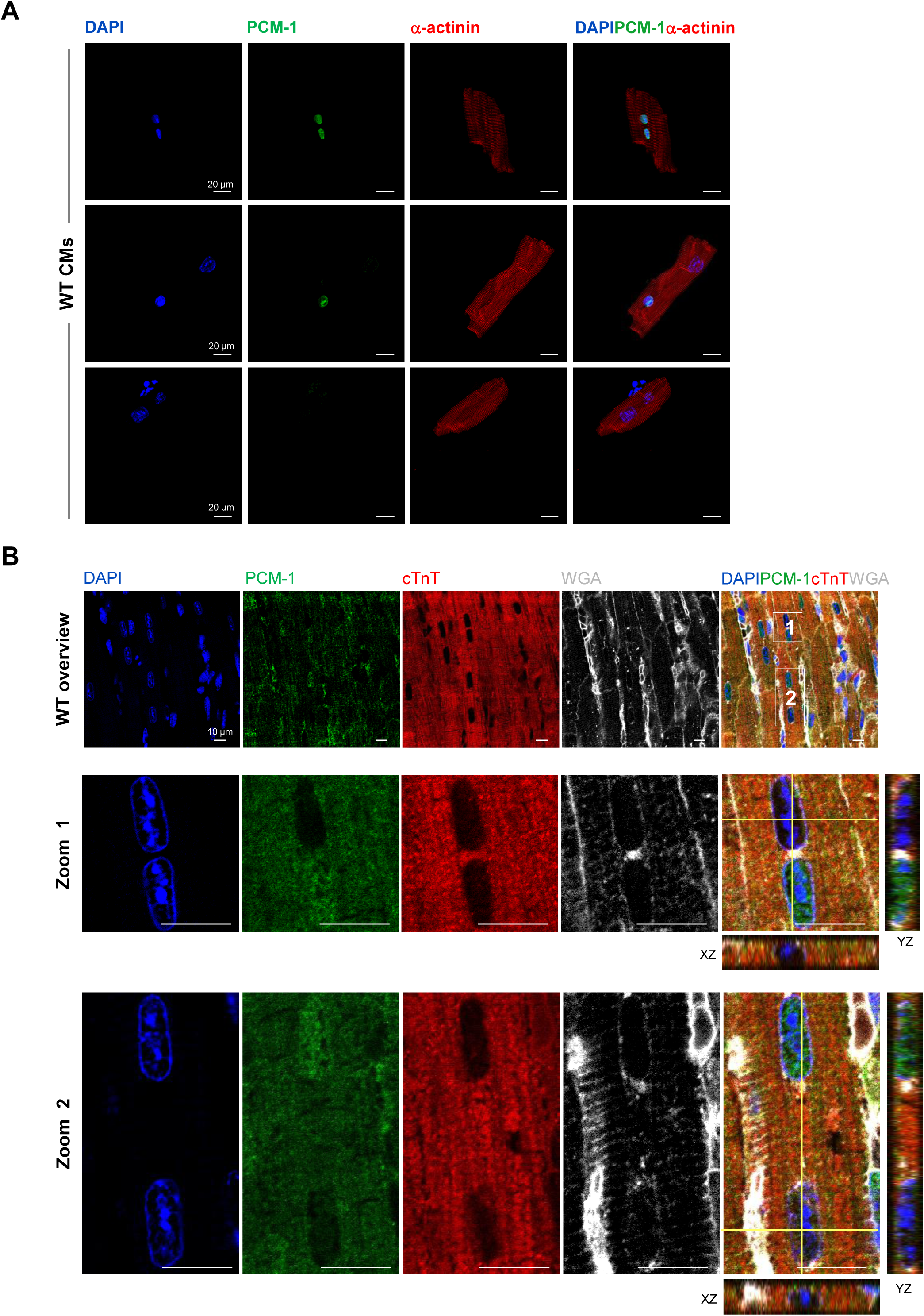
PCM1 as a qualitative but specific CM nuclear marker. **A, B**, PCM1 expression in CMs evaluated (**A**) *in vitro* in isolated CMs (α-actinin^+^ cells) or (**B**) *in situ* (cTnT^+^/WGA cells) in paraffin-embedded heart sections from 2-month old WT mice by immunofluorescence confocal imaging. As shown, although PCM1 was specific for CM nucleus staining, it does not label all CM nuclei both *in vitro* and *in situ*, thus in agreement with the use of PCM1 as a more qualitative but specific CM nuclear marker to confirm resident CM proliferation.

**Figure S5.**
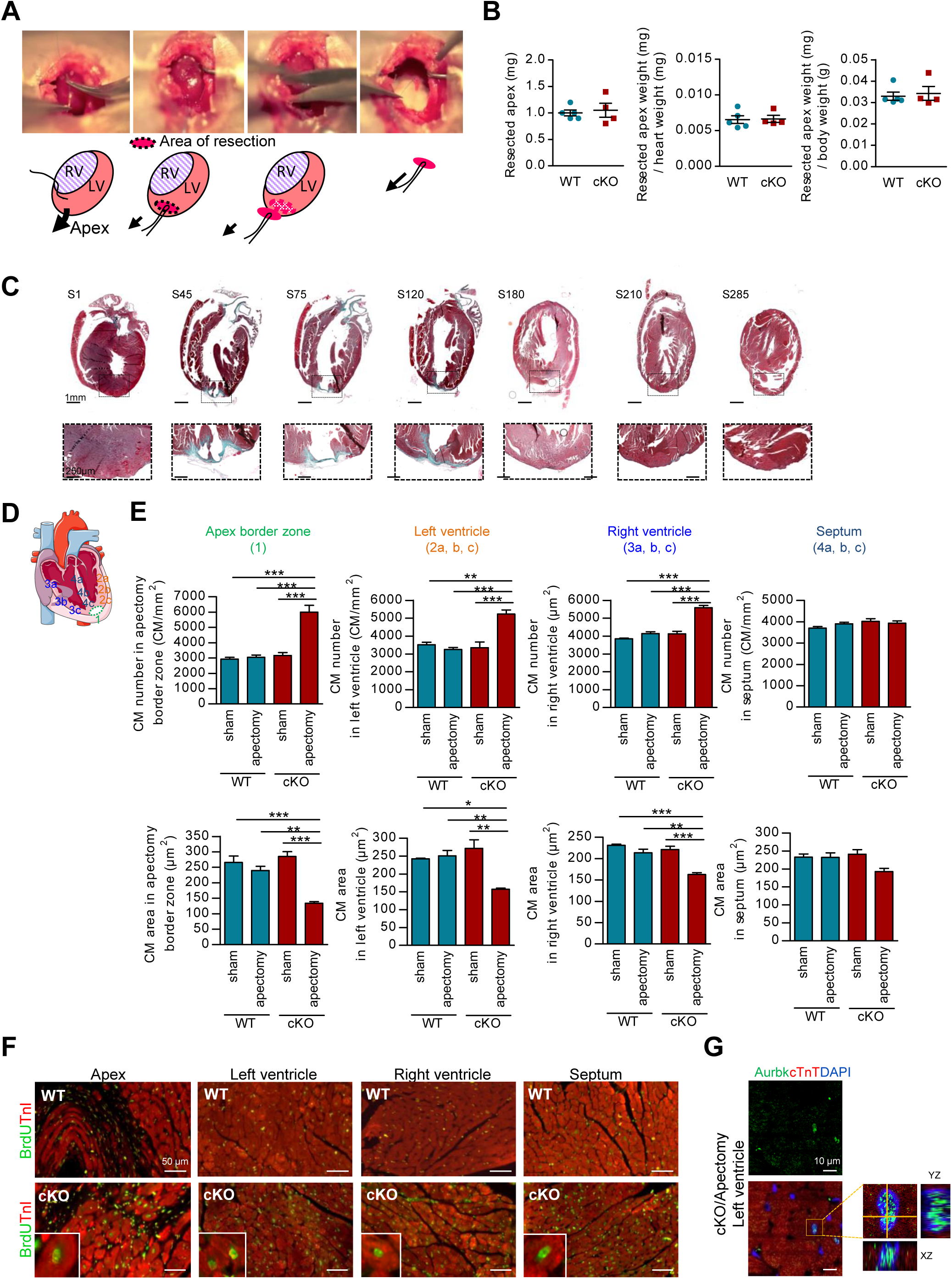
Ephrin-B1 deletion promotes overall cardiac regeneration following apectomy via adult CM proliferation. **A**, Surgical procedure illustration of the apical resection of the adult mouse heart. **B**, Apex weights relative to heart or body weights following resection in hearts from 2-month-old WT or cKO mice (n=4 to 5mice/group). **C**, Scar extent in the apex visualized by Masson’s trichrome staining of serial paraffin-embedded longitudinal sections (5 µm each) from 21 days post-apectomy WT hearts. Sections (S) were performed on the whole heart and only sections delimiting the scar extent are presented. **D**, Illustration of the different heart area quantified in (**E). E**, CM density and area quantified from WGA-stained heart longitudinal cross-sections obtained from apectomized or non-apectomized (sham) 2-month-old WT and *efnb1*^*-/-*^ KO mice 21 days after apectomy. Analysis represent quantification of 3 sections from 3-6 hearts/group. **F-G**, Representative immunofluorescent images of (**F**) CM replication (BrdU^+^/Troponin I, TnI^+^) and (**G**) mitosis (Aurkb^+^/cTnT^+^/DAPI^+^) in the indicated heart compartments of WT and cKO mice 21 days post-apectomy. All data are mean ± SEM.; Student’s *t*-test for 2 group comparisons or one-way ANOVA with Tukey post-hoc test for 4 group comparisons * *P*<0.05, ** *P*<0.01, *** *P*<0.001.

**Figure S6.**
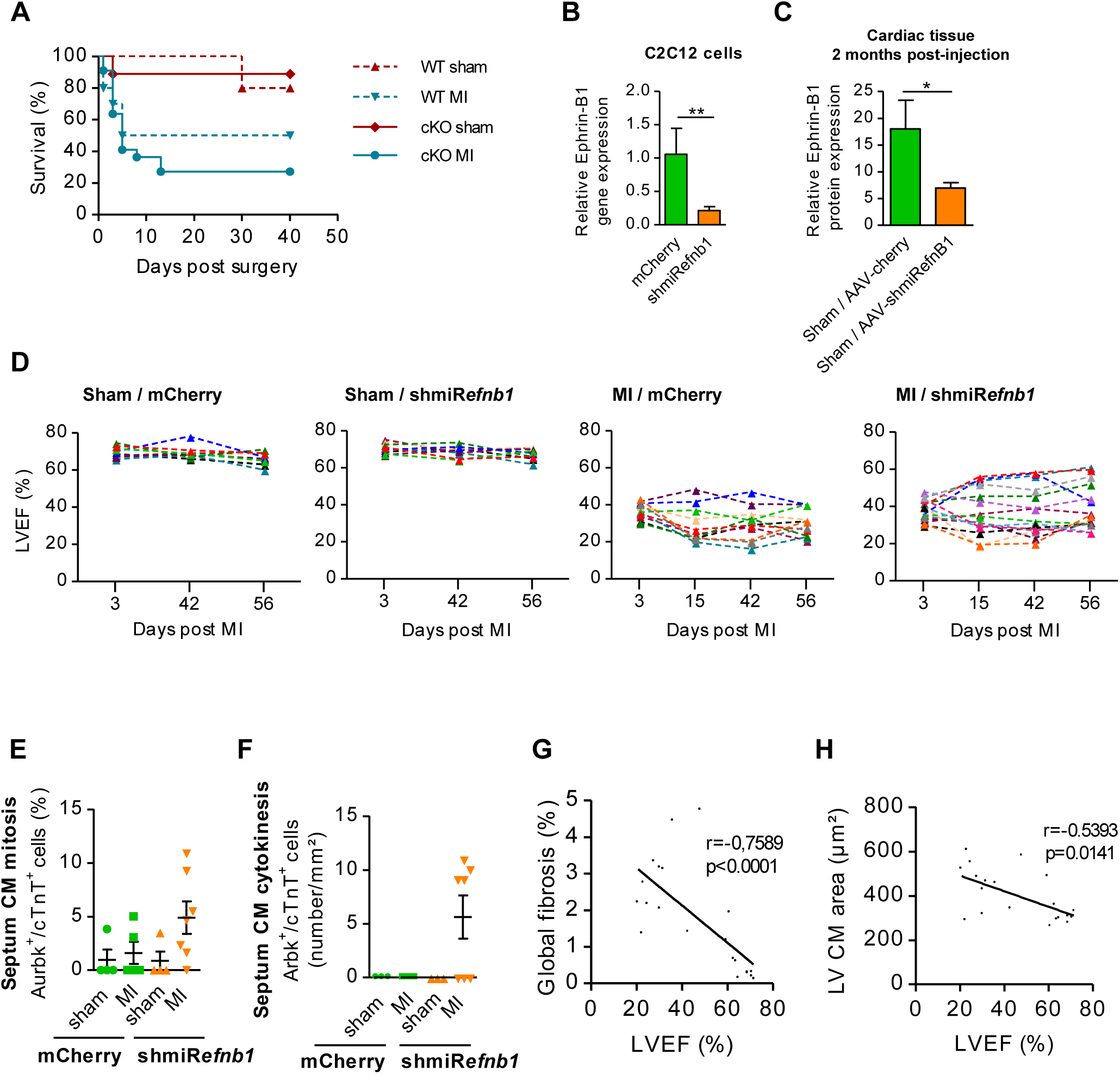
Validation of AAV9-shmir*efnb1* strategy in post-myocardial infarction to evaluate the impact of *efnb1* deletion in cardiac regeneration. **A**, Survival kinetics of 2-month-old WT or cKO mice subjected to left anterior descending coronary artery ligation (Myocardial Infarction, MI) (n=10 to 16 mice/group, 56 days post-MI). **B-C**, Pre-validation of shmiR*efnb1* (pAAV-EPHB147) efficacy (**B**) *in vitro* by qRT-PCR in Ephrin-B1-endogenously-expressing-C2C12 cells (n=4 for each group) and validation (**C**) *in vivo* (AAV9-shmiR*efnb1*) by Western-blot on whole cardiac tissue extract from C57Bl/6J mice 56 days after AAV9-shmiR*efnb1* or control vector (AAV9-mCherry) injections (n=4 mice/group). **D**, Kinetics of Left Ventricle Ejection Fraction (LVEF) from individual mouse in each group (n=10 to 16 mice/group). **E, F**, *In situ* quantifications of CM (**E**) mitosis (Aurkb^+^/DAPI^+^/cTnT^+^) and (**F**) cytokinesis (Aurkb^+^ cleavage furrows/cTnT^+^) in the left ventricle septum (n=4 to 7 mice/group). **G, H**, Ejection fraction correlations with **(G)** global cardiac fibrosis (n=21 mice) or **(H)** left ventricle CM area (n=20 mice). All data are mean ± SEM. Statistical significance was assessed using Student’s *t*-test for 2 group comparisons or one-way ANOVA with Tukey post-hoc test for 4 group comparisons while statistical significance of linear correlation was assessed using Spearman test. * *P*<0.05, ** *P*<0.01, *** *P*<0.001, ns, not significant.

**Figure S7.**
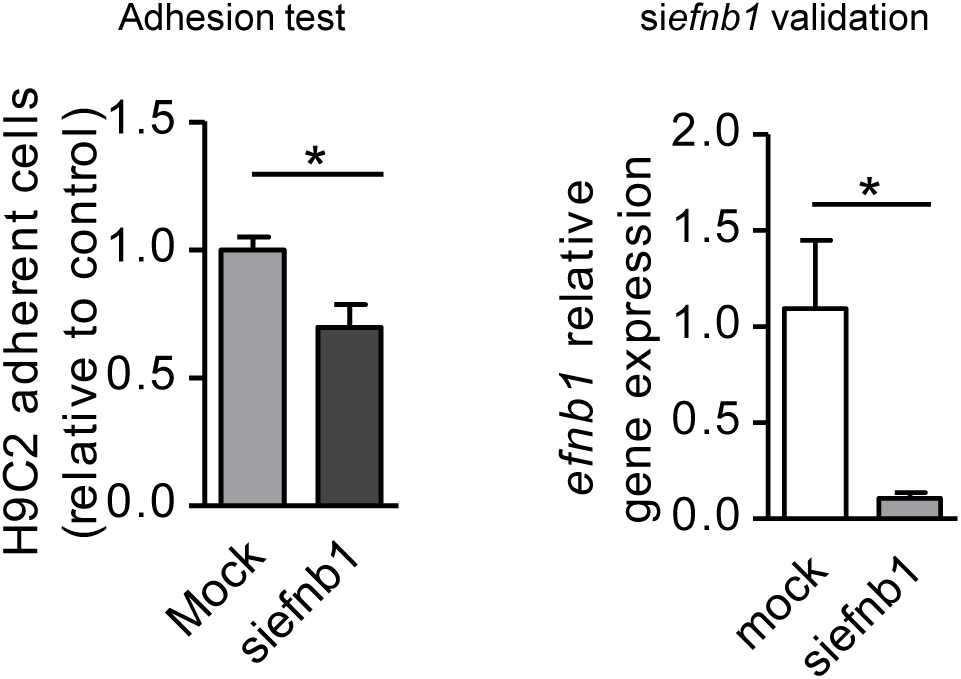
Ephrin-B1 regulates CM adhesion. The role of ephrin-B1 in CM adhesion was evaluated *in vitro* in H9C2 cardiomyoblast cells by siRNA strategy (si*efnb1*) (Left panel) (validation of si*efnb1* efficacy by qRT-PCR in right panel) (n=4 per group).

**Figure S8.**
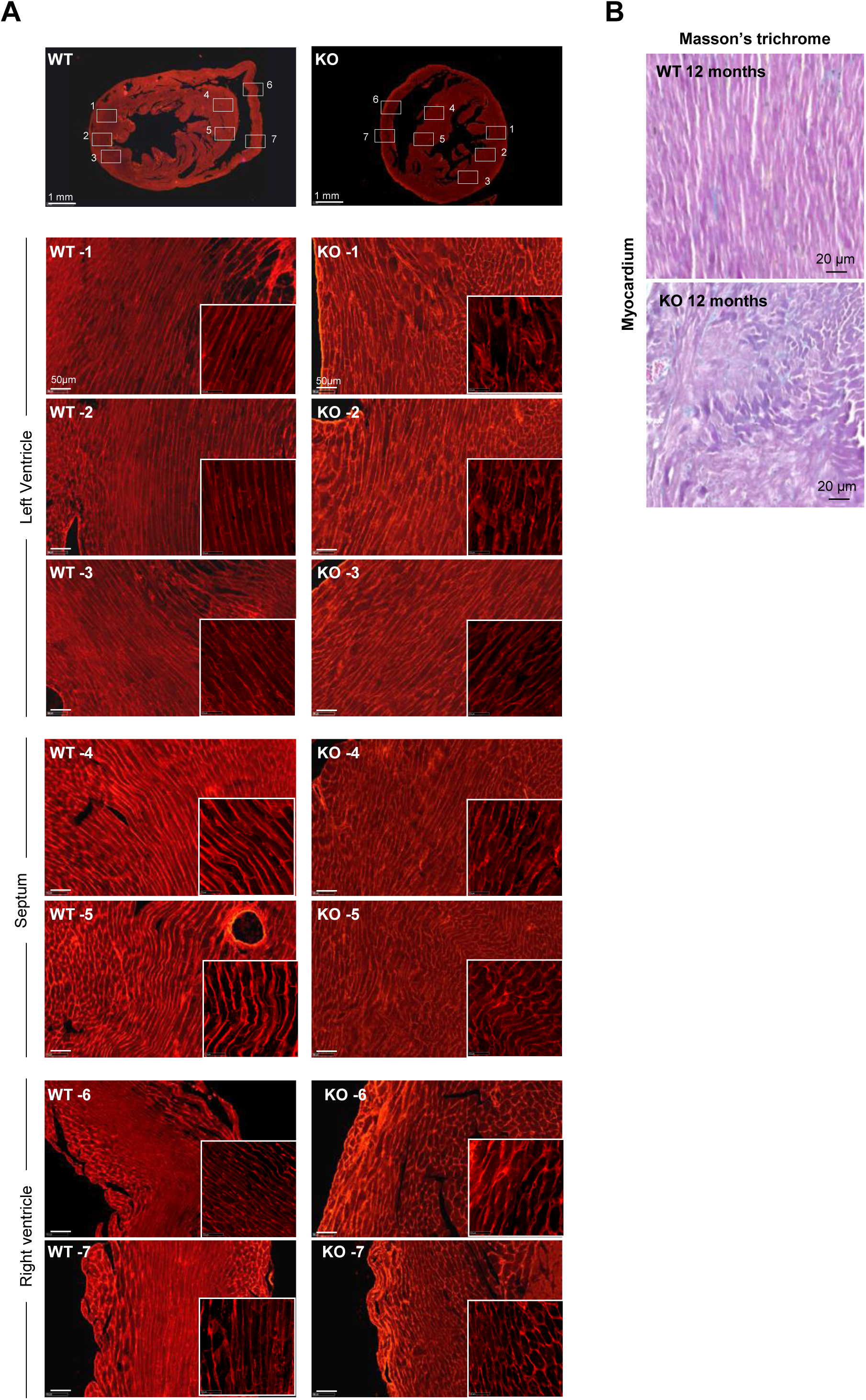
Global disorganization of the cardiac tissue architecture in old-*efnb1*^-/-^ mice. **A**, Wheat-Germ Agglutinin (WGA) staining of heart transversal cross-sections from 12-month-old WT and KO mice showing global disorganization of the cardiac tissue architecture in the whole compartments specifically in the KO mice (LV, Left Ventricle; RV, Right Ventricle). **B**, Representative of Masson’s trichrome staining of cardiac tissue from 12-month-old WT and *efnb1*^*-/-*^ KO mice showing atypical CM rolling in the myocardium in the KO mice.

**Figure S9.**
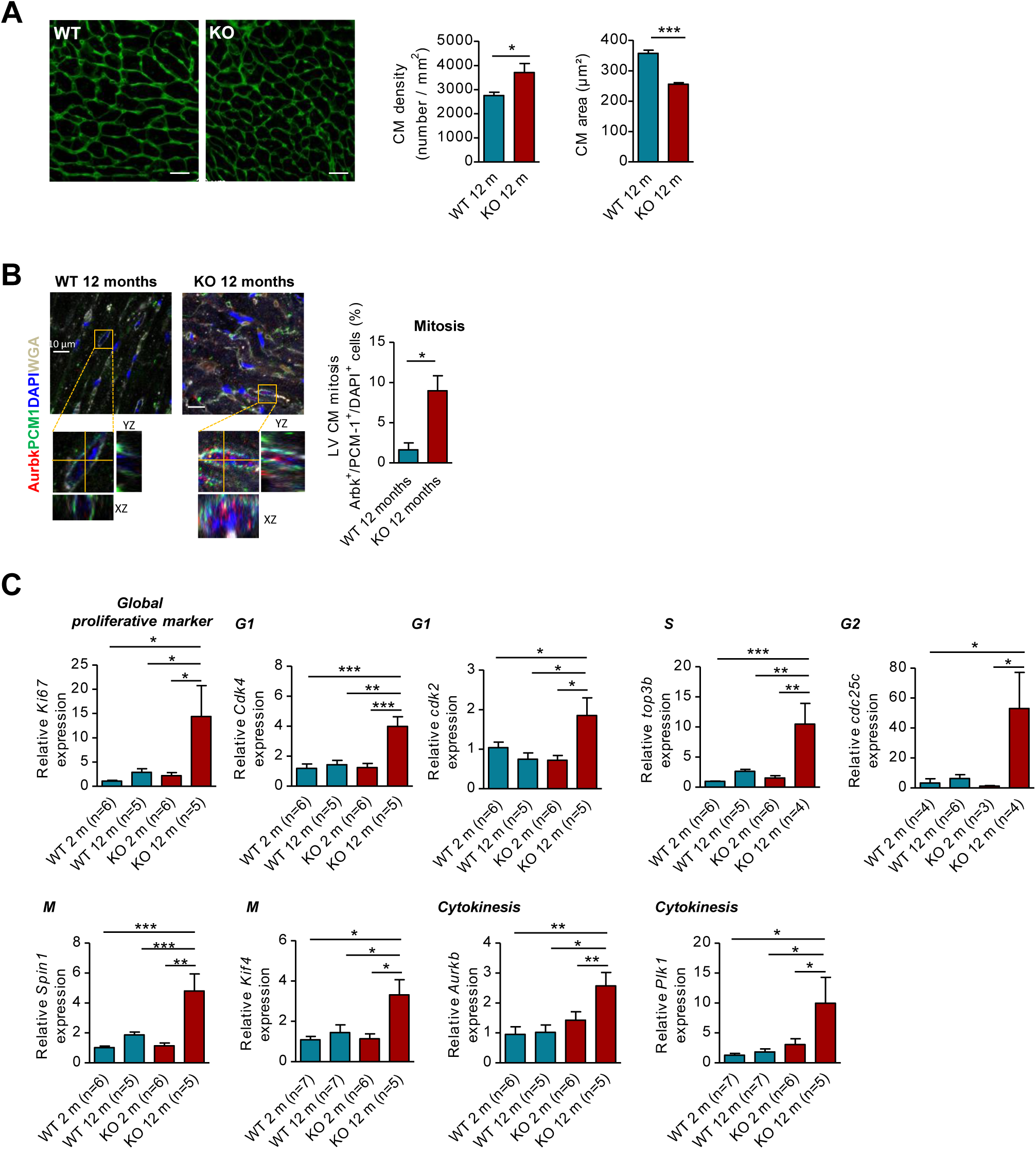
Proliferation indicators of CMs in aged-*efnb1*^-/-^ mice. **A**, CM density and area quantified from WGA-stained heart cross-sections obtained from 12-month-old WT and *efnb1*^*-/-*^ KO mice (∼ 50 CMs/mouse; n=6 mice/group). **B**, *In situ* quantification of resident CM mitosis (PCM1^+^/Aurkb^+^/DAPI^+^) (n=3 mice/group). **C**, qRT-PCR quantification of gene expression of all cell cycle phases (G1, S, G2, M, cytokinesis) (*Cdk4, top3b, cdc25c, Spin1, Aurkb, Ki67, cdk2, Kif4* and *Plk*) in CMs isolated from 2- and 12-month-old WT and *efnb1*^*-/-*^ KO mice. All data are mean ± SEM.; Student’s *t*-test for 2 group comparisons or one-way ANOVA with Tukey post-hoc test for 4 group comparisons * *P*<0.05, ** *P*<0.01, *** *P*<0.001. n.d. not detectable.

**Figure S10.**
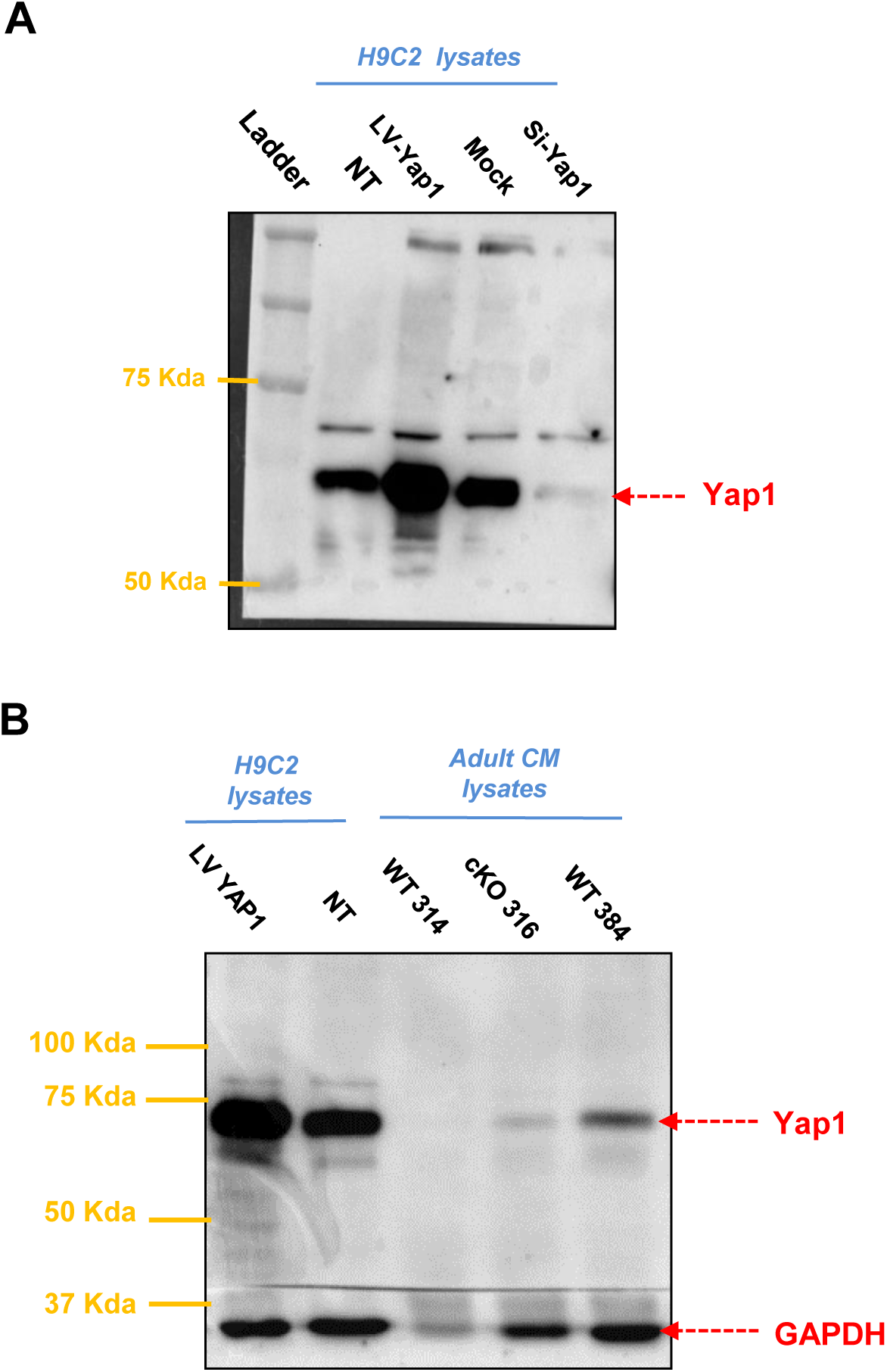
Controls for specific detection of Yap1 in Western-blot. **A**, specificity of Yap1 detection in Western blot was assessed on H9C2 cardiomyoblast cells using cell lysates from H9C2 transduced or not (NT) with a Yap1-encoding lentivirus (LV-Yap1) to overexpress Yap1, or, by contrast, H9C2 transfected with a Si-Yap1 to downregulate Yap1 (si-Yap1). **B**, comparison of cell lysates from H9C2 lysates overexpressing (LV-Yap1) or not (NT) Yap1 with lysates from adult CM isolated from two-month old WT or *efnb1* cKO mice to control for the specificity of Yap1 detection in adult CMs by Western-blot.

**Figure S11.**
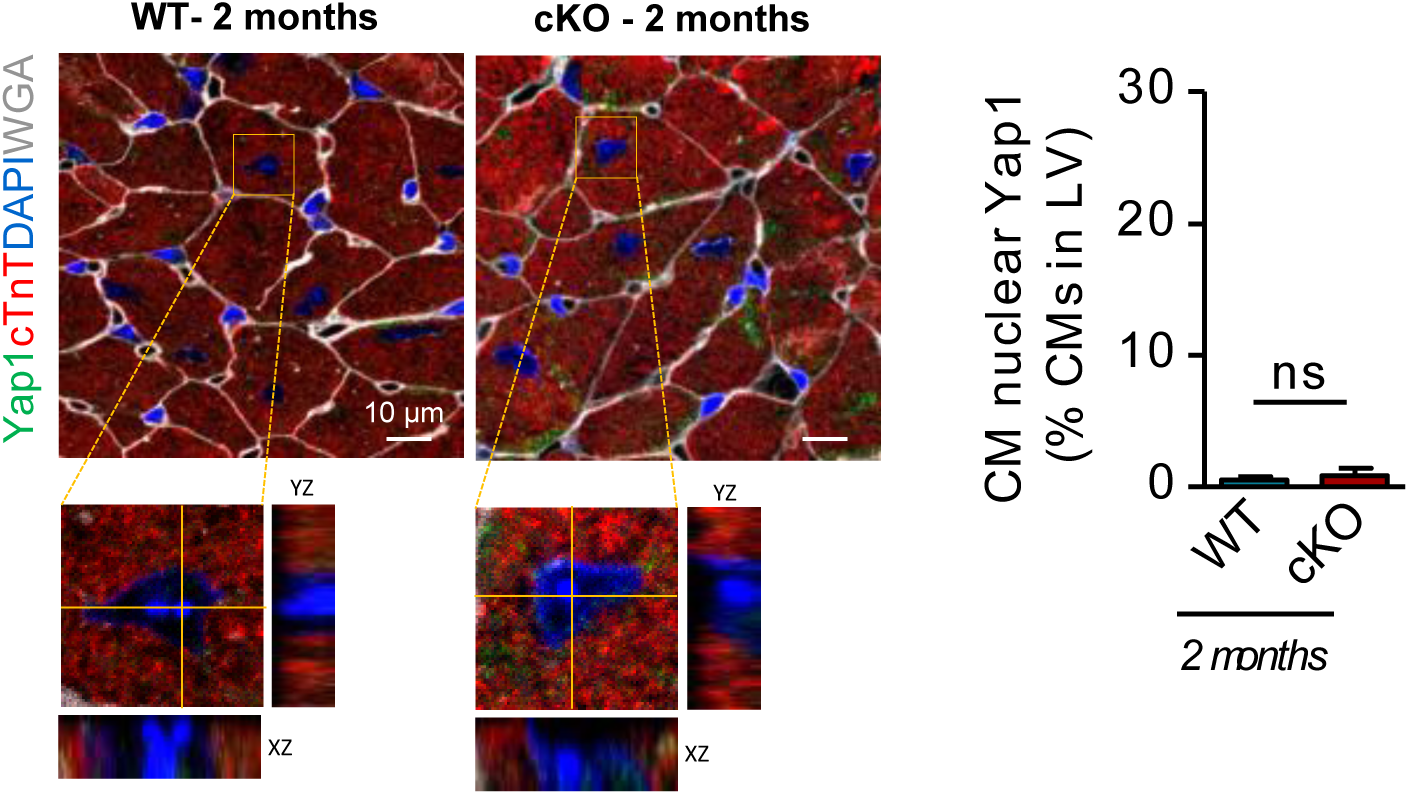
Yap1 is not nuclearized in CMs from resting *efnb1* cKO mice. Activation of Yap-1 in CMs was evaluated by quantification of nuclear Yap1 immunofluorescence staining (Yap1^+^/cTnt^+^/DAPI^+^) in paraffin-sections from 2-month-old WT or cKO mice (n=3 mice/group). Data are expressed as mean ± SEM; Student’s *t*-test. ns, not significant.

**Video S1. Apectomy surgical procedure as described in the detailed method section**.

## References

1. Karra R, Poss KD. Redirecting cardiac growth mechanisms for therapeutic regeneration. J Clin Invest 2017;127(2):427–436.

2. Bergmann O, Bhardwaj RD, Bernard S, Zdunek S, Barnabe-Heider F, Walsh S, Zupicich J, Alkass K, Buchholz BA, Druid H, Jovinge S, Frisen J. Evidence for cardiomyocyte renewal in humans. Science 2009;324(5923):98–102.

3. Kajstura J, Urbanek K, Perl S, Hosoda T, Zheng H, Ogorek B, Ferreira-Martins J, Goichberg P, Rondon-Clavo C, Sanada F, D’Amario D, Rota M, Del Monte F, Orlic D, Tisdale J, Leri A, Anversa P. Cardiomyogenesis in the adult human heart. Circ Res 2010;107(2):305–15.

4. Senyo SE, Steinhauser ML, Pizzimenti CL, Yang VK, Cai L, Wang M, Wu TD, Guerquin-Kern JL, Lechene CP, Lee RT. Mammalian heart renewal by pre-existing cardiomyocytes. Nature 2013;493(7432):433–6.

5. Eulalio A, Mano M, Dal Ferro M, Zentilin L, Sinagra G, Zacchigna S, Giacca M. Functional screening identifies miRNAs inducing cardiac regeneration. Nature 2012;492(7429):376–81.

6. Heallen T, Morikawa Y, Leach J, Tao G, Willerson JT, Johnson RL, Martin JF. Hippo signaling impedes adult heart regeneration. Development 2013;140(23):4683–90.

7. Mahmoud AI, Kocabas F, Muralidhar SA, Kimura W, Koura AS, Thet S, Porrello ER, Sadek HA. Meis1 regulates postnatal cardiomyocyte cell cycle arrest. Nature 2013;497(7448):249–253.

8. Porrello ER, Mahmoud AI, Simpson E, Johnson BA, Grinsfelder D, Canseco D, Mammen PP, Rothermel BA, Olson EN, Sadek HA. Regulation of neonatal and adult mammalian heart regeneration by the miR-15 family. Proc Natl Acad Sci U S A 2013;110(1):187–92.

9. Tian Y, Liu Y, Wang T, Zhou N, Kong J, Chen L, Snitow M, Morley M, Li D, Petrenko N, Zhou S, Lu M, Gao E, Koch WJ, Stewart KM, Morrisey EE. A microRNA-Hippo pathway that promotes cardiomyocyte proliferation and cardiac regeneration in mice. Sci Transl Med 2015;7(279):279ra38.

10. Bassat E, Mutlak YE, Genzelinakh A, Shadrin IY, Baruch Umansky K, Yifa O, Kain D, Rajchman D, Leach J, Riabov Bassat D, Udi Y, Sarig R, Sagi I, Martin JF, Bursac N, Cohen S, Tzahor E. The extracellular matrix protein agrin promotes heart regeneration in mice. Nature 2017;547(7662):179–184.

11. Bersell K, Arab S, Haring B, Kuhn B. Neuregulin1/ErbB4 signaling induces cardiomyocyte proliferation and repair of heart injury. Cell 2009;138(2):257–70.

12. Patterson M, Barske L, Van Handel B, Rau CD, Gan P, Sharma A, Parikh S, Denholtz M, Huang Y, Yamaguchi Y, Shen H, Allayee H, Crump JG, Force TI, Lien CL, Makita T, Lusis AJ, Kumar SR, Sucov HM. Frequency of mononuclear diploid cardiomyocytes underlies natural variation in heart regeneration. Nat Genet 2017;49(9):1346–1353.

13. Leach JP, Heallen T, Zhang M, Rahmani M, Morikawa Y, Hill MC, Segura A, Willerson JT, Martin JF. Hippo pathway deficiency reverses systolic heart failure after infarction. Nature 2017;550(7675):260–264.

14. Li J, Gao E, Vite A, Yi R, Gomez L, Goossens S, van Roy F, Radice GL. Alpha-catenins control cardiomyocyte proliferation by regulating Yap activity. Circ Res 2015;116(1):70–9.

15. Mohamed TMA, Ang YS, Radzinsky E, Zhou P, Huang Y, Elfenbein A, Foley A, Magnitsky S, Srivastava D. Regulation of Cell Cycle to Stimulate Adult Cardiomyocyte Proliferation and Cardiac Regeneration. Cell 2018.

16. Wei K, Serpooshan V, Hurtado C, Diez-Cunado M, Zhao M, Maruyama S, Zhu W, Fajardo G, Noseda M, Nakamura K, Tian X, Liu Q, Wang A, Matsuura Y, Bushway P, Cai W, Savchenko A, Mahmoudi M, Schneider MD, van den Hoff MJ, Butte MJ, Yang PC, Walsh K, Zhou B, Bernstein D, Mercola M, Ruiz-Lozano P. Epicardial FSTL1 reconstitution regenerates the adult mammalian heart. Nature 2015;525(7570):479–85.

17. Porrello ER, Mahmoud AI, Simpson E, Hill JA, Richardson JA, Olson EN, Sadek HA. Transient regenerative potential of the neonatal mouse heart. Science 2011;331(6020):1078–80.

18. Pasumarthi KB, Field LJ. Cardiomyocyte cell cycle regulation. Circ Res 2002;90(10):1044–54.

19. Huang W, Feng Y, Liang J, Yu H, Wang C, Wang B, Wang M, Jiang L, Meng W, Cai W, Medvedovic M, Chen J, Paul C, Davidson WS, Sadayappan S, Stambrook PJ, Yu XY, Wang Y. Loss of microRNA-128 promotes cardiomyocyte proliferation and heart regeneration. Nat Commun 2018;9(1):700.

20. Vite A, Zhang C, Yi R, Emms S, Radice GL. alpha-Catenin-dependent cytoskeletal tension controls Yap activity in the heart. Development 2018;145(5).

21. Anversa P, Olivetti G, Loud AV. Morphometric study of early postnatal development in the left and right ventricular myocardium of the rat. I. Hypertrophy, hyperplasia, and binucleation of myocytes. Circ Res 1980;46(4):495–502.

22. Kullander K, Klein R. Mechanisms and functions of Eph and ephrin signalling. Nat Rev Mol Cell Biol 2002;3(7):475–86.

23. Genet G, Guilbeau-Frugier C, Honton B, Dague E, Schneider MD, Coatrieux C, Calise D, Cardin C, Nieto C, Payre B, Dubroca C, Marck P, Heymes C, Dubrac A, Arvanitis D, Despas F, Altie MF, Seguelas MH, Delisle MB, Davy A, Senard JM, Pathak A, Gales C. Ephrin-B1 is a novel specific component of the lateral membrane of the cardiomyocyte and is essential for the stability of cardiac tissue architecture cohesion. Circ Res 2012;110(5):688–700.

24. Ikenishi A, Okayama H, Iwamoto N, Yoshitome S, Tane S, Nakamura K, Obayashi T, Hayashi T, Takeuchi T. Cell cycle regulation in mouse heart during embryonic and postnatal stages. Dev Growth Differ 2012;54(8):731–8.

25. Poolman RA, Gilchrist R, Brooks G. Cell cycle profiles and expressions of p21CIP1 AND P27KIP1 during myocyte development. Int J Cardiol 1998;67(2):133–42.

26. Sdek P, Zhao P, Wang Y, Huang CJ, Ko CY, Butler PC, Weiss JN, Maclellan WR. Rb and p130 control cell cycle gene silencing to maintain the postmitotic phenotype in cardiac myocytes. J Cell Biol 2011;194(3):407–23.

27. Becker RO, Chapin S, Sherry R. Regeneration of the ventricular myocardium in amphibians. Nature 1974;248(5444):145–7.

28. Mahmoud AI, Porrello ER, Kimura W, Olson EN, Sadek HA. Surgical models for cardiac regeneration in neonatal mice. Nat Protoc 2014;9(2):305–11.

29. Trumble DR, McGregor WE, Kerckhoffs RC, Waldman LK. Cardiac assist with a twist: apical torsion as a means to improve failing heart function. J Biomech Eng 2011;133(10):101003.

30. Bujak M, Kweon HJ, Chatila K, Li N, Taffet G, Frangogiannis NG. Aging-related defects are associated with adverse cardiac remodeling in a mouse model of reperfused myocardial infarction. J Am Coll Cardiol 2008;51(14):1384–92.

31. Walsh S, Ponten A, Fleischmann BK, Jovinge S. Cardiomyocyte cell cycle control and growth estimation in vivo--an analysis based on cardiomyocyte nuclei. Cardiovasc Res 2010;86(3):365–73.

32. Zhang Y, Del Re DP. A growing role for the Hippo signaling pathway in the heart. J Mol Med (Berl) 2017;95(5):465–472.

33. Hong W, Guan KL. The YAP and TAZ transcription co-activators: key downstream effectors of the mammalian Hippo pathway. Semin Cell Dev Biol 2012;23(7):785–93.

34. Mohamed TMA, Ang YS, Radzinsky E, Zhou P, Huang Y, Elfenbein A, Foley A, Magnitsky S, Srivastava D. Regulation of Cell Cycle to Stimulate Adult Cardiomyocyte Proliferation and Cardiac Regeneration. Cell 2018;173(1):104–116 e12.

35. Lin Z, Pu WT. Harnessing Hippo in the heart: Hippo/Yap signaling and applications to heart regeneration and rejuvenation. Stem Cell Res 2014;13(3 Pt B):571–81.

36. Xin M, Olson EN, Bassel-Duby R. Mending broken hearts: cardiac development as a basis for adult heart regeneration and repair. Nat Rev Mol Cell Biol 2013;14(8):529–41.

37. Ragni CV, Diguet N, Le Garrec JF, Novotova M, Resende TP, Pop S, Charon N, Guillemot L, Kitasato L, Badouel C, Dufour A, Olivo-Marin JC, Trouve A, McNeill H, Meilhac SM. Amotl1 mediates sequestration of the Hippo effector Yap1 downstream of Fat4 to restrict heart growth. Nat Commun 2017;8:14582.

38. Morikawa Y, Heallen T, Leach J, Xiao Y, Martin JF. Dystrophin-glycoprotein complex sequesters Yap to inhibit cardiomyocyte proliferation. Nature 2017;547(7662):227–231.

39. Fulford A, Tapon N, Ribeiro PS. Upstairs, downstairs: spatial regulation of Hippo signalling. Curr Opin Cell Biol 2017;51:22–32.

40. Meng Z, Moroishi T, Guan KL. Mechanisms of Hippo pathway regulation. Genes Dev 2016;30(1):1–17.

41. Daar IO. Non-SH2/PDZ reverse signaling by ephrins. Semin Cell Dev Biol 2012;23(1):65–74.

42. Varelas X. The Hippo pathway effectors TAZ and YAP in development, homeostasis and disease. Development 2014;141(8):1614–26.

43. Jopling C, Sleep E, Raya M, Marti M, Raya A, Izpisua Belmonte JC. Zebrafish heart regeneration occurs by cardiomyocyte de-differentiation and proliferation. Nature 2010;464(7288):606–9.

44. Brette F, Luxan G, Cros C, Dixey H, Wilson C, Shiels HA. Characterization of isolated ventricular myocytes from adult zebrafish (Danio rerio). Biochem Biophys Res Commun 2008;374(1):143–6.

## Supplementary References

1. Genet G, et al. (2012) Ephrin-B1 is a novel specific component of the lateral membrane of the cardiomyocyte and is essential for the stability of cardiac tissue architecture cohesion. Circ Res 110(5):688–700.

2. Bergmann O & Jovinge S (2012) Isolation of cardiomyocyte nuclei from post-mortem tissue. J Vis Exp (65).

3. Leone M, Magadum A, & Engel FB (2015) Cardiomyocyte proliferation in cardiac development and regeneration: a guide to methodologies and interpretations. Am J Physiol Heart Circ Physiol 309(8):H1237–1250.

4. Bruel A & Nyengaard JR (2005) Design-based stereological estimation of the total number of cardiac myocytes in histological sections. Basic Res Cardiol 100(4):311–319.

5. Doevendans PA, Daemen MJ, de Muinck ED, & Smits JF (1998) Cardiovascular phenotyping in mice. Cardiovasc Res 39(1):34–49.

6. Reliene R & Schiestl RH (2006) Differences in animal housing facilities and diet may affect study outcomes-a plea for inclusion of such information in publications. DNA Repair (Amst) 5(6):651–653.

7. Alkass K, et al. (2015) No Evidence for Cardiomyocyte Number Expansion in Preadolescent Mice. Cell 163(4):1026–1036.

8. Prats AC, et al. (2013) CXCL4L1-fibstatin cooperation inhibits tumor angiogenesis, lymphangiogenesis and metastasis. Microvasc Res 89:25–33.

9. Bellot M, et al. (2015) Dual agonist occupancy of AT1-R-alpha2C-AR heterodimers results in atypical Gs-PKA signaling. Nat Chem Biol 11(4):271–279.

